# Protective function and differentiation cues of brain-resident CD8+ T cells during immune surveillance of chronic latent *Toxoplasma gondii* infection

**DOI:** 10.1101/2024.02.08.579453

**Authors:** Rémi Porte, Marcy Belloy, Alexis Audibert, Emilie Bassot, Amel Aïda, Marine Alis, Romain Miranda-Capet, Aurélie Jourdes, Klaas van Gisbergen, Frédérick Masson, Nicolas Blanchard

## Abstract

Chronic *T. gondii* infection induces brain-resident CD8+ T cells (bTr) but their protective functions and differentiation cues remain undefined. Here, we used a mouse model of latent infection by *T. gondii* leading to effective CD8+ T cell-mediated parasite control. Thanks to antibody depletion approaches, we found that peripheral circulating CD8+ T cells are dispensable for brain parasite control during chronic stage, indicating that CD8+ bTr are sufficient to prevent brain parasite reactivation. We observed that the retention markers CD69, CD49a and CD103 are sequentially acquired by brain parasite-specific CD8+ T cells throughout infection, and that a majority of CD69/CD49a/CD103 triple-positive (TP) CD8+ T cells also express Hobit, a transcription factor associated with tissue residency. This TP subset develops in a CD4+ T cell-dependent manner, and is associated with effective parasite control during chronic stage. Conditional invalidation of TAP-mediated MHC class I presentation showed that presentation of parasite antigens by glutamatergic neurons and microglia regulate the differentiation of CD8+ bTr into TP cells. Single-cell transcriptomic analyses upon *T. gondii* latency vs. encephalitis revealed that resistance to encephalitis is associated with the expansion of stem-like subsets of CD8+ bTr.

In summary, parasite-specific brain-resident CD8+ T cells are functionally heterogeneous and autonomously ensure parasite control during *T. gondii* latent infection. Their differentiation is shaped by neuronal and microglial MHC I presentation. A more detailed understanding of local T cell-mediated immune surveillance of this common parasite is needed for harnessing brain-resident CD8+ T cells in order to enhance control of chronic brain infections.

## Introduction

CD8+ T cells are key players of adaptive immune responses against infections, both through their ability to kill cells infected by an intracellular pathogen and through their capacity to produce cytokines (e.g. IFN-γ) that activate anti-pathogen responses in neighbouring cells. Upon cognate antigen recognition, costimulation and cytokine signaling, naïve CD8+ T cells may differentiate into short-lived effectors cells (SLEC) that secrete effector molecules (e.g. IFN-γ and granzyme B), or into memory precursor cells (MPC) (Belz and Kallies, 2010). MPC can give rise to 3 types of memory cells: (i) central memory cells (Tcm) that have stem-like properties and remain mostly within lymphoid organs, where they are poised for being recalled, (ii) effector memory cells (Tem) that navigate between peripheral tissues and lymphoid organs, and are equipped to quickly produce cytokines and cytotoxic factors upon restimulation, (iii) resident memory cells (Trm) that durably station within non-lymphoid tissues, including the central nervous system (CNS), where they provide local protective immunity against re-infection (Gebhardt et al., 2009; Heeg and Goldrath, 2023; Steinert et al., 2015; van Gisbergen et al., 2021).

These principles have been established mostly based on settings in which the pathogen is eventually cleared. The persistence of a pathogen beyond acute infection can profoundly perturb, or at the very least complexify, this scenario. A famous illustration is the functional impairment (“exhaustion”) observed in a fraction of virus-specific CD8+ T cells during systemic chronic viral infections such as HIV (Day et al., 2006) or LCMV (Wherry et al., 2003) infections (Lan et al., 2023). How CD8+ Trm differentiate in the context of tissue-restricted latent infections has been partially addressed with HSV-2 in the skin and sensory ganglia (Park et al., 2023; Roychoudhury et al., 2020), or with Polyoma virus in the brain (Mockus et al., 2018; Ren et al., 2020). Yet, the extent by which the Trm compartment is shaped by the long-term cohabitation with pathogens that remain transcriptionally, and “antigenically”, active throughout chronic infection (e.g. as in *Toxoplasma gondii* infection), remains poorly understood.

Generally, CD8+ Trm do not express a single defining marker. Depending on the pathophysiologial context and tissue of residency, CD8+ Trm may upregulate all or a combination of surface molecules involved in tissue retention, such as the C-type lectin CD69 (encoded by the Cd69 gene), the α1 integrin CD49a (Itga1 gene), and the αE integrin CD103 (Itgae gene). Typically, Trm also display higher expression of transcription factors that simultaneously facilitate expression of the tissue residency program and suppress circulatory-associated genes, such as RUNX3 (Runx3 gene) (Milner et al., 2017; Zitti et al., 2023), BLIMP1 (Prdm1 gene) (Kragten et al., 2018), and HOBIT (Zfp683 gene) (Mackay et al., 2016).

Although the requirements are expected to differ according to the tissue microenvironment (Heeg and Goldrath, 2023), several general mechanisms underlie the differentiation and maintenance of CD8+ Trm across tissues. They include local antigen recognition and TCR signal strength (Abdelbary et al., 2023; Low et al., 2020; Maru et al., 2017; Sanecka et al., 2018), engagement of costimulatory receptors such as ICOS (Peng et al., 2022), signaling by cytokines such as type I IFN and IL-12 (Bergsbaken et al., 2017), IL-15 (Mackay et al., 2015), TGF-β (Ferreira et al., 2020; Mackay et al., 2015; Mani et al., 2019) and IL-21 (Ren et al., 2020; Tkachev et al., 2021), the latter illustrating the key contribution of CD4+ T cell help (Frieser et al., 2022; Son et al., 2021; Vincenti et al., 2022) in optimal CD8+ Trm differentiation. These cues are thought to induce a network of factors, chiefly transcription factors, that both control expression of surface adhesion molecules and orchestrate the metabolic adaptations of Trm to their specific site of residence (Reina-Campos et al., 2023). This explains the phenotypical heterogeneity of Trm across different tissues. Interestingly, even within a given tissue, an important diversity has been observed, potentially mirrorring the different functional subsets that comprise the circulating memory CD8+ T cell compartment (Heeg and Goldrath, 2023; Milner et al., 2020; Park et al., 2023) (reviewed in (Konjar et al., 2022; Szabo, 2023)).

To this date, the extent of CD8+ Trm functional diversity, its underlying mechanisms and long-term functional consequences have remained ill-defined in settings of tissue-restricted chronic infections. To fill this gap, we have set-out to study the function, differentiation cues and heterogeneity of brain-resident memory CD8+ T cells (bTrm) cells in the context of chronic cerebral infection by the brain-persisting *Toxoplasma gondii* (*T. gondii*) parasite.

*T. gondii* is a foodborne, obligate intracellular, apicomplexan parasite, considered as the most widespread and generalist zoonotic pathogen on earth as it is found at every latitude and is able to infect all warm-blooded animals. Following infection, rapidly dividing *T. gondii* tachyzoites can invade all nucleated cells and disseminate systemically within the host, inducing strong type I immune responses. Several parasite effectors are secreted into host cells during acute infection, thereby eliciting immune modulatory mechanisms and preventing clearance of the parasite (Hakimi et al., 2017). When inside retinal, muscular or neuronal cells, tachyzoites may convert into bradyzoites, which are contained within intracellular cysts. Encysted bradyzoite stages are responsible for the long-term, chronic persistence of *T. gondii* in the CNS, a phase called latency. Based on the ∼30% worldwide seroprevalence of *T. gondii* in humans (Bigna et al., 2020), and on the fact that ∼2/3 of these exposed individuals display immunoglobulins specific for latent *T. gondii* stages (Dard et al., 2021), it is estimated that more than 1.5 billion humans carry *T. gondii* cysts in the brain. Beside the risk of congenital malformations which may occur in case of primary infection during pregnancy, and beside the severe forms of toxoplasmosis caused by hypervirulent South American strains of *T. gondii*, the most severe clinical manifestations of *T. gondii* infection are usually restricted to individuals presenting with acquired immune suppression, with a specific association with HIV/AIDS. In individuals previously exposed to *T. gondii*, defective cellular immune surveillance resulting from AIDS may lead to parasite reactivation (i.e. bradyzoite to tachyzoite reverse conversion) and uncontrolled tachyzoite replication, ultimately causing systemic disseminated toxoplasmosis and/or, most frequently, *T. gondii* encephalitis (TE). TE clinical symptoms combine headache, fever, neurological deficits and epileptic manifestations. Before the introduction of effective antiretroviral therapy against HIV, TE was the most frequent inaugural manifestation of HIV/AIDS in Europe. More than 13 million people worldwide are currently estimated to be co-infected with HIV and *T. gondii*, with nearly 90% of them in sub-Saharan Africa, making TE the second infectious cause of mortality among people living with HIV in Africa (Wang et al., 2017). Therefore, TE is a global public health challenge that imposes a major burden on Africa (Vidal, 2019). Notably, although the links remain mostly correlative for now (Johnson and Koshy, 2020; Laing et al., 2020), mounting evidence suggests that *T. gondii* contributes to neuropsychological disorders in immunocompetent individuals (Burgdorf et al., 2019), and that it worsens cognitive decline upon aging (Bayani et al., 2019; Nimgaonkar et al., 2016).

Up to now, most of the knowledge generated about immune surveillance of *T. gondii* in the brain, is based on mouse models. Resident neuroimmune cells such as microglia and astrocytes play an important role, for example through IL-1α production by microglia (Batista et al., 2020), and IL-33 and CCL2 release by astrocytes, which are important to mobilize other immune cells (Orchanian et al., 2024; Still et al., 2020). Innate immune cells recruited from the periphery, such as monocytes (Biswas et al., 2015) and type 1 innate lymphoid cells (ILC1) (Steffen et al., 2022), are also implicated, with ILC1 being an early source of IFN-γ in response to cerebral infection. IFN-γ is indeed a critical cytokine for *T. gondii* restriction in the CNS. Accordingly, STAT1-mediated signaling in astrocytes (Hidano et al., 2016) and microglia (Cowan et al., 2022) is required for optimal parasite containment in the brain.

In part owing to their ability to produce IFN-γ, CD4+ (i.e. Th1) and CD8+ T lymphocytes are considered as major players in *T. gondii* restriction. This is well established in the context of acute infection (Nishiyama et al., 2020), which aligns with the observation that perforin, a key component of CD8+ T cell cytotoxicity appears dispensable during acute infection (Denkers et al., 1997). The respective contributions of IFN-γ and cytotoxic factors are not yet fully clarified during chronic infection but perforin-dependent CD8+ T cell cytotoxicity was proposed to play a prominent role in limiting brain parasite burden (Denkers et al., 1997; Suzuki et al., 2010). In agreement, studies comparing mouse strains that are differentially susceptible to TE indicated that CD8+ T cells are a pillar of protective immunity during chronic infection. Efficient presentation of parasite-derived antigens by molecules of the class I Major Histocompatibility Complex (MHC) (Blanchard et al., 2008; Brown et al., 1995; Feliu et al., 2013), and more specifically MHC I presentation of tachyzoite-derived antigens by infected, glutamatergic neurons of the CNS, drives effective parasite control in the early stages of parasite brain invasion, resulting in low cyst burden at chronic stage (Salvioni et al., 2019). Two pioneer studies (one in an encephalitis-susceptible (Landrith et al., 2017) and one in an encephalitis-resistant (Sanecka et al., 2018) model) have reported the presence of brain CD8+ T cells with a surface phenotype (e.g. CD103+) and a transcriptional profile that are consistent with tissue residency. A subsequent study confirmed the presence in the brain of parasite-specific CD69+ CD103+ CD8+ T cells, which display a Trm signature and had been recently activated (Shallberg et al., 2022). Active parasite replication was reported to have a limited impact on the persistence of parasite-specific CD69+ CD103+ CD8+ bTrm during *T. gondii* encephalitis, suggesting that the phenotype of such cells may be imprinted early post-brain invasion (Shallberg et al., 2022). Yet, as of now, little is known about the protective role, functional diversity and differentiation cues of parasite-specific CD8+ bTrm during *T. gondii* latency. Notably, the roles played by brain-specific antigen-presenting cells such as neurons and microglia, remain unaddressed.

In this work, we have set-out to address the function, differentiation cues and transcriptional heterogeneity of CD8+ bTrm using mouse models of chronic *T. gondii* infection. We have accumulated evidence that brain-resident CD8+ T cells play a more prominent role than brain-circulating CD8+ T cells in keeping parasites in check during latency. We also report that both excitatory neurons and CX3CR1+ brain macrophages (including microglia), optimize the differentiation of the brain-resident CD8+ T cell compartment. At last, by delineating the transcriptional heterogeneity of parasite-specific CD8+ bTrm upon encephalitis and throughout early *vs*. late latency, we identify stem-like T cell populations that are preferentially associated with long-term parasite control.

## Results

Latency is considered as the most frequent outcome in immunocompetent individuals exposed to *T. gondii*. Upon infection with type II *T. gondii* strains, the susceptible C57BL/6 mice develop TE. Hence, to be able to study *T. gondii*-specific CD8+ bTrm throughout latent infection, we infected C57BL/6 mice with a *T. gondii* line derived from the type II Prugnaud (Pru) strain, modified to express a model antigen (GRA6-OVA(Feliu et al., 2013; Salvioni et al., 2019)) that is efficiently processed and presented by MHC class I molecules, thereby eliciting TE-protective CD8+ T cell responses in the brain (Salvioni et al., 2019)) and leading to latent infection (**Fig. 1A**).

**Figure 1.**
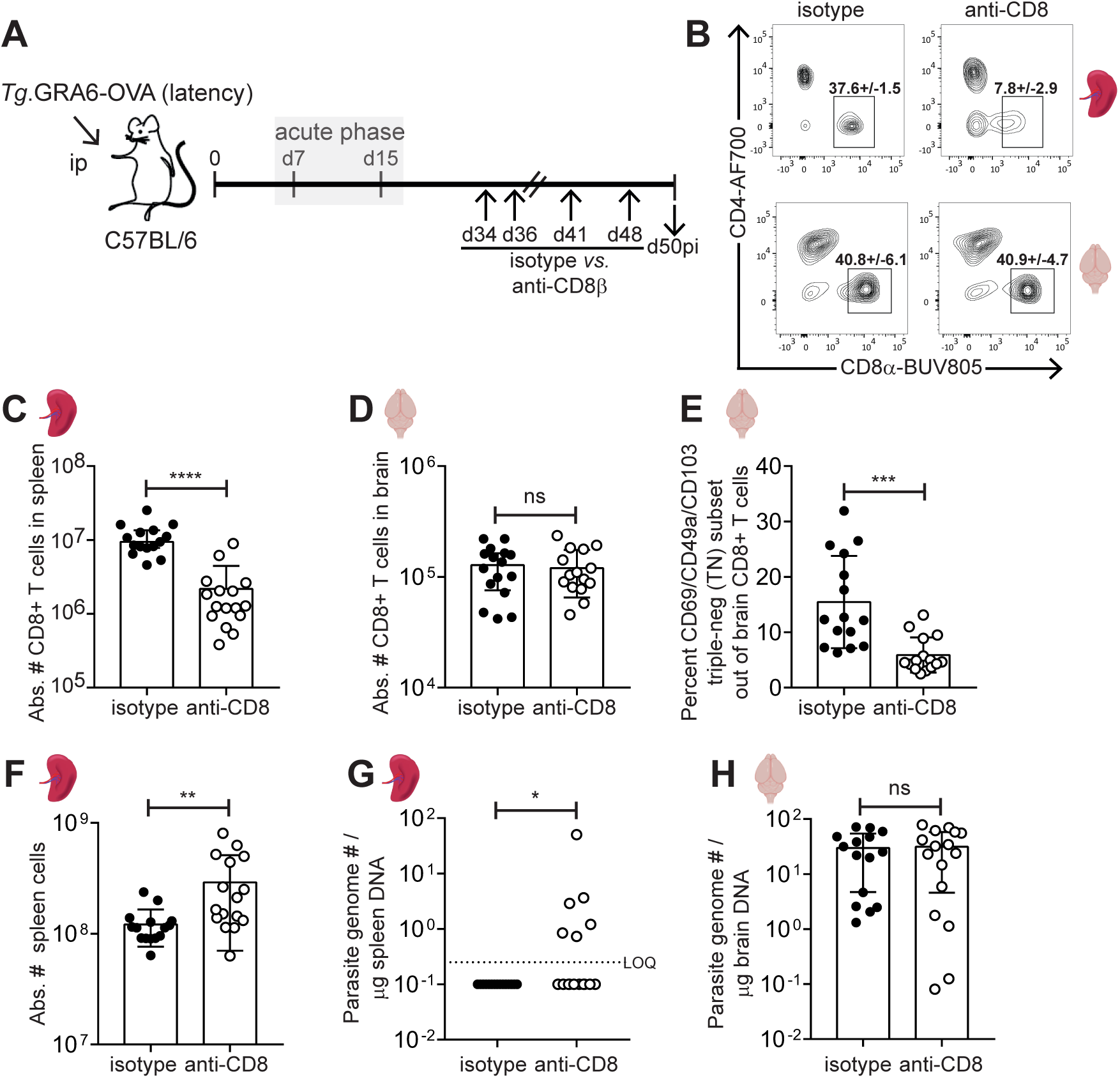
Peripheral CD8+ T cell depletion during latent *T. gondii* infection leads to recrudescence of parasite in spleen but not in brain. (A) Schematics of experimental workflow: C57BL/6 mice were infected with 200 tachyzoites of GRA6-OVA-expressing *T. gondii* Pru. At chronic phase (d34pi), mice were administered with an anti-CD8β antibody or an isotype control, twice at d34pi & d36pi, and then once a week for 2 weeks until d50pi. (B) Representative Facs plots showing effects of anti-CD8β treatment in spleen and brain. Numbers on plots show the percentage +/- s.d of CD8+ T cells out of single, live, CD3+ T cells. (C, D) Bar graphs showing absolute number of CD8+ T cells in spleen (C) and brain (D) at d50pi. (E) Bar graph showing the proportion of CD69/CD49a/CD103 triple-negative subset (circulating) among brain CD8+ T cells at d50pi. (F) Bar graph showing total number of spleen cells at d50pi, reflecting the splenomegaly associated with anti-CD8 depletion. (C, D, E, F) Each dot represents one mouse. Bars show the mean ± SD of N = 15 (isotype) *vs*. 16 (anti-CD8β) mice per group, pooled from 3 independent experiments. Mann-Whitney tests between isotype-treated and anti-CD8-treated groups. (G) Spleen parasite burden measured by qPCR on genomic DNA extracted from spleen. Each dot represents one mouse with N = 15 (isotype) *vs*. 16 (anti-CD8β) mice per group, pooled from 3 independent experiments. Dotted line indicates the limit of quantification. Parasite DNA was detectable in 6 out of 16 anti-CD8β -treated mice, *vs*. 0 out of 15 isotype-treated control mice (p=0.0177, Fisher exact test). (H) Brain parasite burden measured by qPCR on genomic DNA extracted from brain. Each dot represents one mouse with N = 15 (isotype) *vs*. 16 (anti-CD8β) mice per group, pooled from 3 independent experiments. (C, F) Mann-Whitney test between isotype-treated and anti-CD8β-treated groups. (D, E, H) Unpaired t-test between isotype-treated and anti-CD8β-treated groups.

To first evaluate the respective contributions of circulating *vs*. brain-resident CD8+ T cells (bTr) in immune surveillance of *T. gondii* infection during chronic stage, we took advantage of the differential sensitivity of circulating and bTr to the depletion induced by systemic administration of an anti-CD8 antibody. Such a treatment was indeed reported to preferentially eliminate peripheral circulating CD8+ T cells while largely preserving brain-resident T cells (Frieser et al., 2022; Steinbach et al., 2016). Starting at ∼1 month post-infection, we administered an anti-CD8β antibody for 2 weeks, (**Fig. 1A**). This treatment depleted CD8+ T cells in the spleen (**Fig. 1B, 1C**) but, as expected, it had a limited effect on CD8+ T cells from the brain (**Fig. 1B, 1D**). The only subset of CD8+ T cells that was significantly reduced in the brain was the CD69/CD49a/CD103 triple-negative (TN) population, which are most likely circulating cells (**Fig. 1E**). Two (2) weeks post-treatment, the anti-CD8 antibody treatment led to splenomegaly (not shown) and to an increase in total splenocyte number (**Fig. 1F**), potentially reflecting a higher parasite burden in the spleen. A recrudescence of *T. gondii* in the spleen was indeed observed in ∼40% (6 out of 16) treated animals (**Fig. 1G**). However, this anti-CD8 treatment did not alter parasite control in the brain (**Fig. 1H**), indicating that peripheral CD8+ T cells are important for parasite control in the spleen but dispensable for parasite control in the brain during latency, in turn suggesting that an anti-CD8-insensitive, brain-resident, CD8+ T cell compartment is self-sufficient to limit reactivation during chronic stage.

In order to monitor the formation of brain-resident CD8+ T cells throughout infection in this model, we isolated CD8+ T cells from the brain at different times post-infection (**Fig. 2A**), and used flow cytometry to measure the surface expression of 3 tissue retention molecules (CD69, CD49a and CD103) on both parasite (OVA)-specific CD8α+ T cells and total CD8α+ T cells (**Fig. 2B, 2C**). At both chronic infection timepoints examined (d32pi and d76pi), the majority of total and parasite-specific CD8+ T cells in the brain co-expressed CD69 and CD49a (i.e. belonged to the DP or TP subsets) (**Fig. 2D, 2E**), indicative of their likelihood to reside in the brain and, based on CD49a expression, suggestive of a type 1/cytotoxic profile (Cheuk et al., 2017; Park et al., 2023). CD103 was largely absent in the acute stage (d13pi) and was observed exclusively on cells already expressing CD69 and CD49a. Accordingly, while few parasite-specific CD8+ T cells simultaneously expressed the 3 retention markers at d13pi, almost half of them acquired triple expression of CD69, CD49a and CD103 (TP subset) at d32pi, a proportion which remained stable until d76pi (**Fig. 2E**). Notably, during chronic stage, the proportion of TP cells was slightly higher among parasite (OVA)-specific CD8+ T cells than among total CD8+ T cells, but overall, the kinetic evolution of these 3 markers was similar for parasite-specific and total CD8+ T cells (**Fig. 2D, 2E**). Using a tdTomato-based reporter mouse for Hobit (Homolog of Blimp-1 in T cells), a transcription factor typically associated with tissue residency in T cells (Mackay et al., 2016; Parga-Vidal et al., 2021) (**Fig. 3A**), we observed that CD69+ CD49a+ CD8+ T cells, positive or not for CD103 (i.e. TP and DP subsets respectively), contain the highest proportion of Hobit-tdTomato+ cells (**Fig. 3B, 3C**), showing that, in these settings, Hobit+ brain-resident CD8+ bTrm comprise CD103-negative and CD103-positive cells. On this basis, we inferred (i) that most DP and TP cells are *bona fide* (Hobit+) brain-resident T cells and (ii) that most TN cells (i.e. CD69/CD49a/CD103 triple-negative) are T cells that are recirculating in the brain. Considering the persistence of their cognate antigen throughout chronic stage, the ‘memory’ status of such OVA-specific resident T cells may be questioned. Thus, we preferred to name them brain-resident T cells (bTr) instead of brain-resident memory T cells (bTrm). Interestingly, compared to TN bTcirc, DP/TP bTr displayed a higher percentage of ‘bi-functional’ cells able to co-express IFN-γ and granzyme B upon PMA/ionomycin stimulation (**Fig. 3D-F)**, showing that DP/TP bTr are endowed with a higher effector potential than CD8+ bTcirc, further reinforcing the notion that DP/TP bTr play an important role in brain parasite restriction.

**Figure 2.**
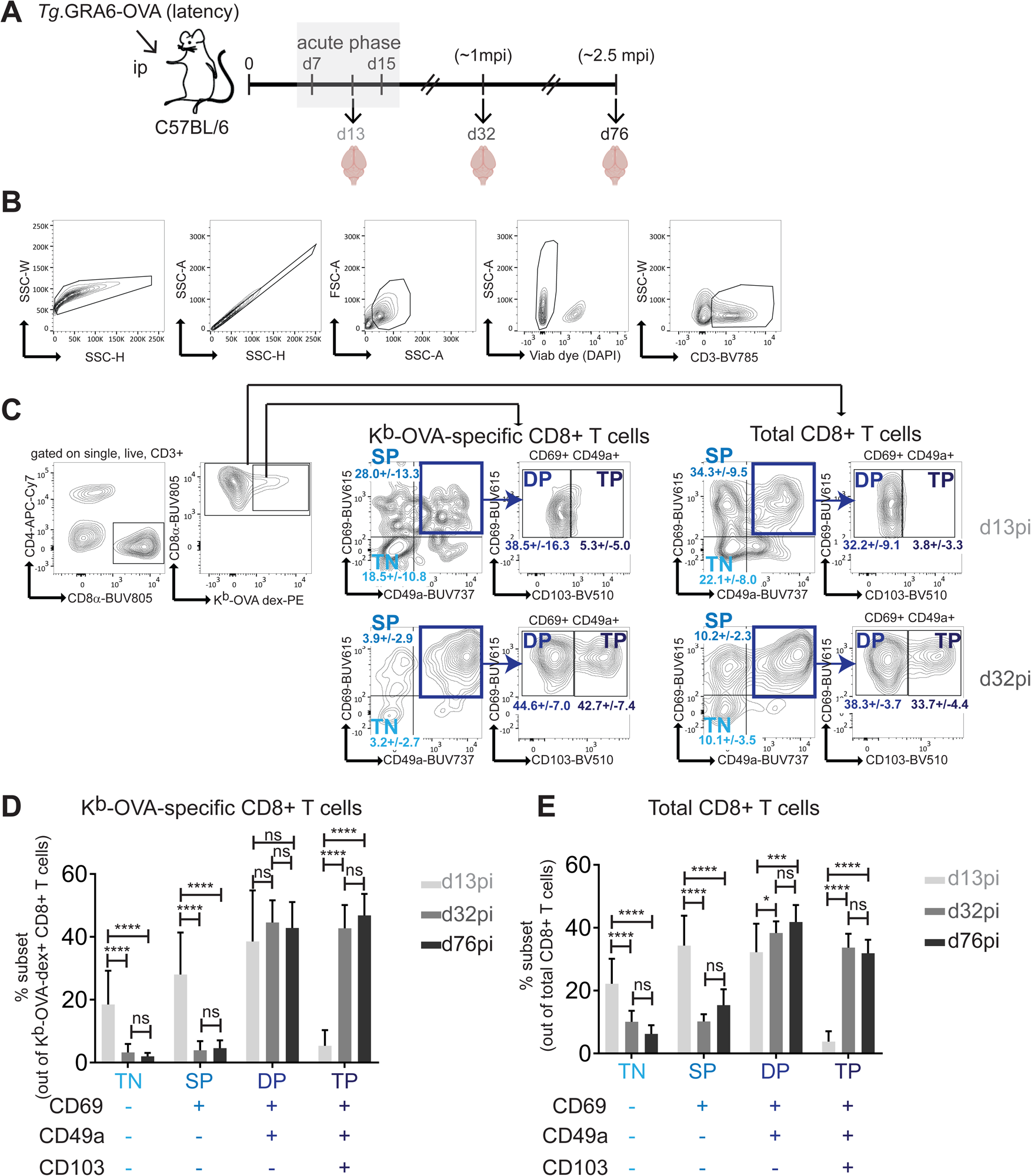
Kinetics of CD69, CD49a and CD103 surface expression on brain-isolated CD8α+ T cells during *T. gondii* infection. (A) Schematics of experimental workflow: C57BL/6 mice were infected intra-peritoneally with 200 tachyzoites of GRA6-OVA-expressing *T. gondii* Pru. Brain-isolated cells were analyzed by flow cytometry at acute (d13pi) and chronic (d32pi and d76pi) stages. (B, C) Gating strategy to analyze surface expression of CD69, CD49a and CD103 on total CD8+ *vs. T. gondii*-specific (dex K^b^-OVA+) CD8+ T cells from the brain by flow cytometry. Numbers on Facs plots show the percentage +/- s.d of each subset (TN: CD69-CD49a-CD103-, SP: CD69+ CD49a-CD103-, DP: CD69+ CD49a+ CD103-, TP: CD69+ CD49a+ CD103+) out of K^b^-OVA-specific CD8+ T cells or out of total CD8+ T cells, as indicated. (D, E) Graphs represent the percentage of each subset out of parasite (OVA)-specific CD8+ T cells (D) or out of total CD8+ T cells (E). Bars show mean ± SD of N = 17 mice at d13pi (pooled from 3 experiments), N = 12 mice at d32pi (pooled from 3 experiments), N = 10 mice at d76pi (from 1 experiment). Two-way ANOVA with Tukey’s multiple comparison test applied on the 3 groups, for every subset.

**Figure 3.**
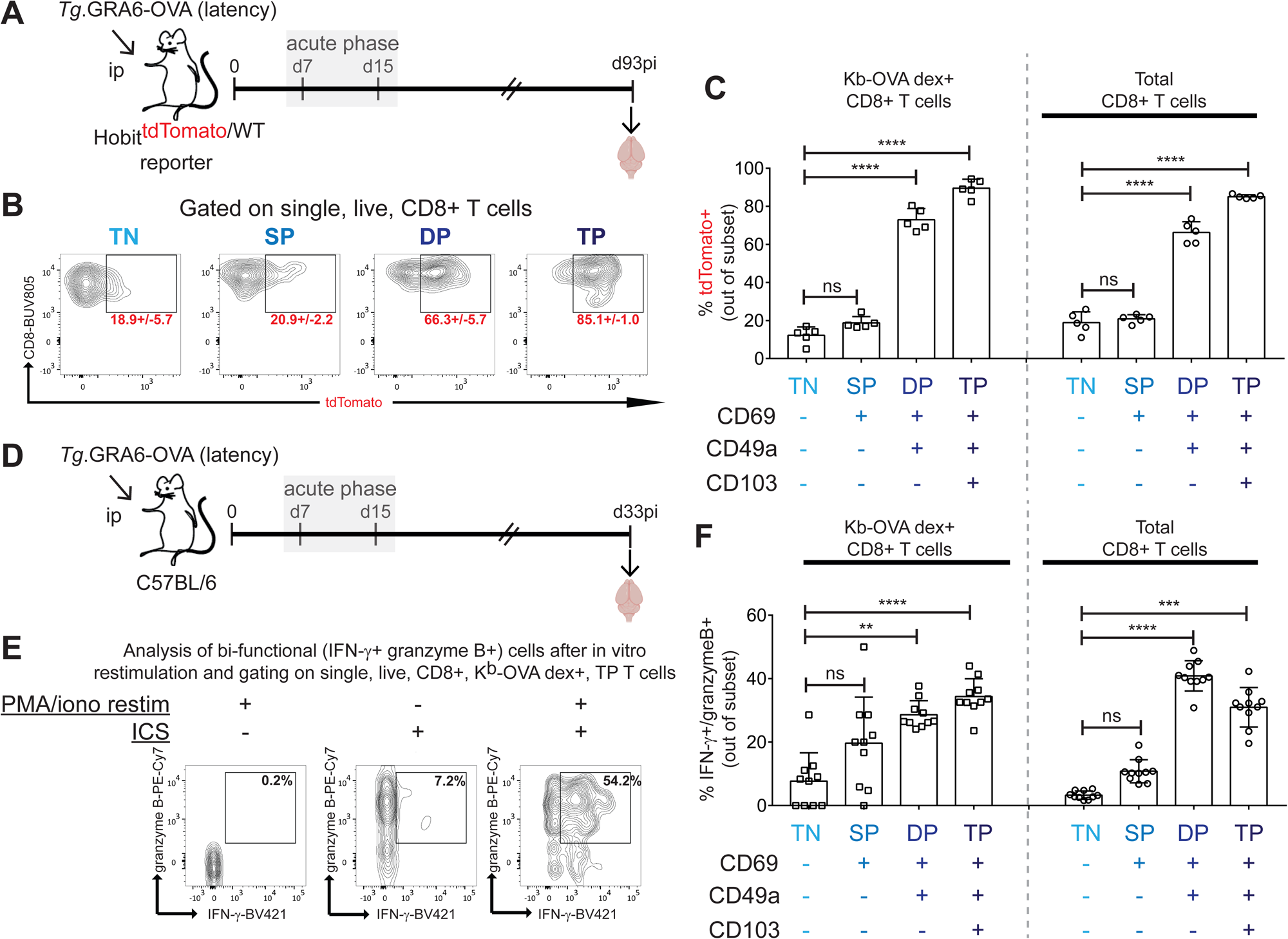
Expression of Hobit, IFN-γ and granzyme B in brain-isolated CD8+ T cells during *T. gondii* latent infection. (A) Schematics of experimental workflow: Hobit-tdTomato reporter mice were infected intra-peritoneally with 200 tachyzoites of GRA6-OVA-expressing *T. gondii* Pru, and analyzed at chronic stage (d93pi). (B) Gating strategy to analyze proportion of Hobit-tdTomato+ cells out of each CD8+ T cell subset (TN: CD69- CD49a- CD103-, SP: CD69+ CD49a- CD103-, DP: CD69+ CD49a+ CD103-, TP: CD69+ CD49a+ CD103+). Red numbers on dot plots show the percentage +/- s.d of tdTomato+ cells out of each CD8+ T cell subset. (C) Graph showing the percentage of tdTomato+ cells out of each subset among parasite (OVA)-specific CD8+ T cells (left part) or among total CD8+ T cells (right part). Bars show mean ± s.d. of N = 5 mice from one experiment, representative of two. Kruskall-Wallis with Dunn’s multiple comparison test used to compare each subset to the TN reference. (D) Schematics of experimental workflow: C57BL/6 mice were infected intra-peritoneally with 200 tachyzoites of GRA6-OVA-expressing *T. gondii* Pru, and analyzed at chronic stage (d33pi). (E) Gating strategy to analyze proportion of cells co-expressing IFN-γ and granzyme B (so-called “bi-functional”) following *ex vivo* PMA/ionomycin restimulation. Shown are representative IFN-γ/granzyme B Facs plots of TP K^b^-OVA dextramer+ CD8+ T cells, restimulated or not with PMA/ionomycin, stained intracellularly or not for IFN-γ and granzyme B, as indicated. Numbers on plots show the percentage of bi-functional cells out of the parental TP K^b^-OVA dextramer+ CD8+ T cells. (F) Graph showing the percentage of bi-functional cells following PMA/ionomycin stimulation and intracellular staining, out of each subset from parasite (OVA)-specific CD8+ T cells (left part) or from total CD8+ T cells (right part). Bars show mean ± s.d. of N = 10 mice from one experiment, representative of 3. Kruskall-Wallis with Dunn’s multiple comparison test used to compare each subset to the TN reference.

To start deciphering the signals driving the formation of CD8+ bTr in this latent infection, we first assessed the importance of cues provided by CD4+ T cells. Hence, we eliminated the CD4+ T cells before *T. gondii* infection with an anti-CD4 depleting antibody and pursued the depletion throughout chronic phase (**Fig. 4A**). As expected, there was a drop in the number of CD4+ T cells in the spleen and brain at acute stage (d13pi) (**Sup. Fig. 1A, 1B, 1D**) but no major effect on the number of parasite (OVA)-specific CD8+ T cells in these 2 organs (**Sup. Fig. 1C, 1E**), and no change in parasite dissemination or access to the brain (**Sup. Fig. 1F, 1G**). This is in line with previous work with CD4 knock-out mice showing that CD4+ T cells are dispensable to resist acute *T. gondii* infection (Schaeffer et al., 2009). At chronic stage (d33pi), the anti-CD4 treatment also reduced the proportion and abundance of CD4+ T cells in the spleen (**Fig. 4B, 4D**) and the brain (**Fig. 4C, 4E**) without negatively impacting the number of parasite (OVA)-specific CD8 T cells in the spleen (**Fig. 4F, 4H**) and brain (**Fig. 4G, 4I**). Strikingly, in the brain, the absence of CD4+ T cells blocked the differentiation of *T. gondii*-specific DP CD8+ T cells into TP cells (**Fig. 4J, 4K**) and reduced the proportion of granzyme B^+^ cells among DP/TP (CD69^+^ CD49a^+^) bTr (**Fig. 4L**). Concomitantly, CD4+ T cell-depleted mice displayed a significantly higher parasite load in the brain (**Fig. 4M**), suggesting that among parasite-specific CD8+ bTr cells, the CD4+ T cell-dependent cytotoxic subsets play a particularly important role in parasite control.

**Figure 4.**
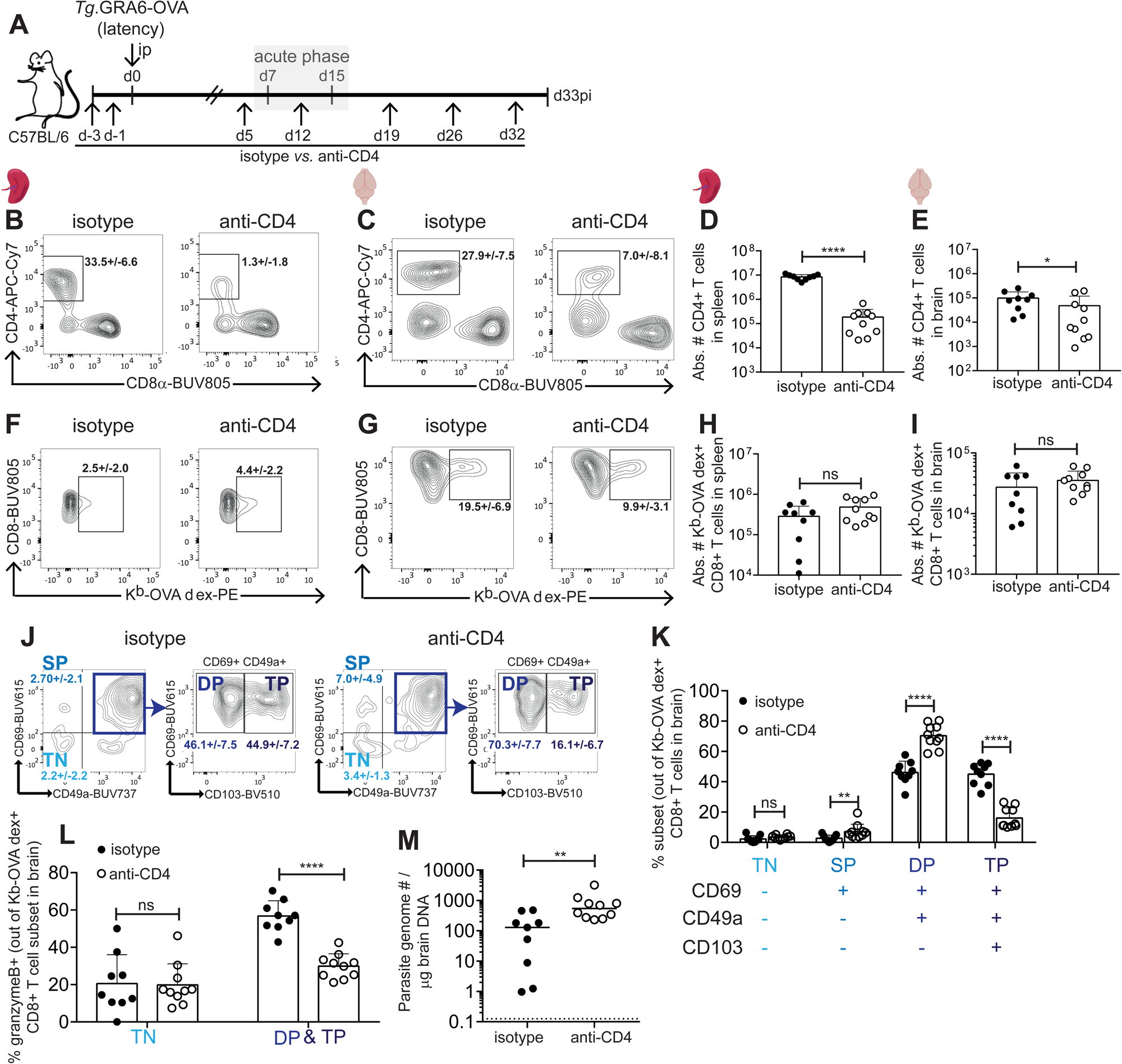
CD4+ T cells drive the differentiation of TP and cytotoxic parasite-specific brain-resident CD8+ T cells, thereby optimizing brain parasite control upon chronic stage. (A) Schematics of experimental workflow: C57BL/6 mice were administered with an anti-CD4 depleting antibody at day -3 and -1 before infection with 200 tachyzoites of GRA6-OVA-expressing *T. gondii* Pru. Injection of anti-CD4 antibody was repeated at d5pi and maintained once per week onwards, until chronic stage. (B, C) Representative contour plots of CD4/CD8 staining after gating on single, live, CD3+ T cells from spleen (B) or brain (C). Numbers on Facs plots show the percentage +/- s.d of CD4+ T cells out of CD3+ T cells. (D, E) Bar graph showing absolute number of CD4+ T cells in spleen (D) and brain (E) at d33pi. (F, G) Representative contour plots of K^b^-OVA dextramer staining after gating on single, live, CD3+ CD8+ T cells from spleen (F) or brain (G). Numbers on dot plots show the percentage +/- s.d of K^b^-OVA dextramer+ T cells out of CD8+ T cells. (H, I) Bar graph showing absolute number of parasite (OVA)-specific CD8+ T cells in spleen (H) and brain (I) at d33pi. (D, E, H, I) Each dot represents one mouse. Bars represent the mean ± SD of N = 9 *vs*. 10 mice per group, pooled from 2 independent experiments with between 4 and 5 mice per group in each experiment. (D, E) Mann-Whitney test between isotype-treated and anti-CD4-treated groups. (H, I) Unpaired t-test between isotype-treated and anti-CD4-treated groups. (J) Representative contour plots showing surface expression of CD69, CD49a and CD103 on OVA-specific (K^b^-OVA dextramer+) CD8+ T cells isolated from the brain. Numbers on Facs plots show the percentage +/- s.d. of each subset (TN: CD69- CD49a- CD103-, SP: CD69+ CD49a- CD103-, DP: CD69+ CD49a+ CD103-, TP: CD69+ CD49a+ CD103+) out of K^b^-OVA-specific CD8+ T cells. (K) Graph showing the percentage of each subset out of parasite (OVA)-specific CD8+ T cells. Bars show mean ± SD of N = 9 *vs*. 10 mice per group, pooled from 2 independent experiments with between 4 and 5 mice per group in each experiment. Mann-Whitney test performed for SP and unpaired t-tests performed for TN, DP, TP between isotype-treated and anti-CD4-treated groups. (L) Bar graph showing the percentage of granzyme B+ cells out of Tcirc (TN, left) or bTr (DP plus TP, right) parasite (OVA)-specific CD8+ T cells. Bars show the mean +/- s.d. of N = 9 *vs*. 10 mice per group, pooled from 2 independent experiments with between 4 and 5 mice per group in each experiment. Mann-Whitney test performed for the TN subset and unpaired t-test performed for the DP+TP subsets, between isotype-treated and anti-CD4-treated group. (M) Brain parasite burden measured by qPCR on genomic DNA extracted from brain. Dotted line indicates the limit of quantification. Line shows mean of N = 9 *vs*. 10 mice per group, pooled from 2 independent experiments with between 4 and 5 mice per group in each experiment. Mann-Whitney test performed to compare isotype-treated and anti-CD4-treated group.

We then wished to gain insights into the local mechanisms that control differentiation of parasite-specific CD8+ bTr cells. Given the relevance of neurons and microglia during brain *T. gondii* infection, we addressed to which extent MHC I presentation of parasite antigens by neurons or microglia, regulates the differentiation of parasite-specific CD8+ bTr. To conditionally disrupt MHC class I processing and presentation, we generated a new Cre/Lox system enabling cell type-specific invalidation of the Tap1 gene, which encodes one subunit of the transporter associated with antigen processing (TAP) (Tap1fl/fl) (**Sup. Fig. 2A**). TAP imports antigenic peptides in the endoplasmic reticulum, where loading of the final antigenic peptides onto nascent MHC class I molecules takes place. Therefore, TAP is a critical component of the MHC I pathway both for endogenous and for exogenous antigens (i.e. cross-presentation) (Adiko et al., 2015; Mantel et al., 2021). In order to target excitatory neurons and brain macrophages (including microglia), we used inducible Cre-reporter systems based on the Camk2a (Casanova et al., 2001) and the Cx3cr1 (Parkhurst et al., 2013) promoters, respectively. To validate the neuron-specific deletion of Tap1, we treated Tap1fl/fl x Camk2a-CreER+ (TAP^neuronKO^) and CreER— (TAP^neuronWT^) mice with tamoxifen, and prepared genomic DNA from neuronal *vs.* non-neuronal cell fractions (**Sup. Fig. 2B**). As expected, the Cre-recombined locus was detectable only in neurons from tamoxifen-treated Cre+ mice (**Sup. Fig. 2C**). Contrary to adult neurons, microglia is amenable for flow cytometry analysis, thus, in order to validate the Tap1fl/fl x Cx3cr1-CreER model, we treated mice with tamoxifen, waited for one month to allow replenishment of the CX3CR1+ monocyte compartment (**Sup. Fig. 2D**), infected the mice and measured surface expression of H-2K^b^ and H-2D^b^ MHC I molecules by Facs on brain-isolated CD45lo CD11b+ cells (comprising microglia and other brain macrophages), CD45hi CD11b+ cells (comprising monocytes) and CD45hi CD11b-(comprising lymphocytes) (**Sup. Fig. 2E**). Expectedly as a consequence of Tap1 invalidation, both the proportion of MHC I-positive cells and the intensity of MHC I staining were drastically reduced in microglia/macrophages from tamoxifen-treated Cre+ animals compared to tamoxifen-treated Cre— mice. This was true for both the H-2K^b^ (**Sup. Fig. 2F-H**) and H-2D^b^ (**Sup. Fig. 2I-K**) alleles of MHC I. Cre induction by tamoxifen had no effect on MHC I expression in the 2 other cell types analyzed (monocytes and lymphocytes) (**Sup. Fig. 2F-K**). For the sake of simplicity, we named Tap1fl/fl x Cx3cr1-Cre+ mice: TAP^microgliaKO^ and Cre— mice: TAP^microgliaWT^. After validating this model, we could evaluate the importance of TAP-dependent MHC I presentation by neurons and microglia in the differentiation of parasite (OVA)-specific CD8+ bTr in the brain. Our previous data had underscored the critical role of neuronal MHC I presentation for parasite control at early chronic stage (Salvioni et al., 2019). We thus anticipated a different parasite burden between TAP^neuronKO^ and TAP^neuronWT^ mice, potentially resulting in different amounts of neuroinflammation and available antigen. To mitigate this confounding factor, we treated mice with an anti-parasitic drug (pyrimethamine) during acute stage, late enough so as to not clear the parasite, but early enough so as to equalize the brain parasite load at d33pi (**Fig.5A, 5B**). We then quantified the percentage (**Fig. 5C**) and absolute number (**Fig. 5D**) of TN, SP, DP and TP subsets among parasite (OVA)-specific CD8+ T cell subsets at 33dpi. TAP invalidation in excitatory neurons led to a slight reduction in the proportion of TP cells and a significant decrease in the abundance of DP & TP parasite (OVA)-specific CD8+ T cells in the brain (**Fig. 5C, 5D**). Moreover, TAP^neuronKO^ mice had less bi-functional IFN-γ+/granzyme B+ OVA-specific bTr (**Fig. 5E**). TAP invalidation in CX3CR1+ microglia/macrophages (**Fig. 5F, 5G**) affected the differentiation of CD8+ bTr as well, mostly also by reducing the TP population of CD8+ bTr (**Fig. 5H, 5I**). However, it did not significantly affect the number of bi-functional IFN-γ+/granzyme B+ OVA-specific CD8+ bTr at d33pi (**Fig. 5J**). In conclusion, presentation of parasite antigens by excitatory neurons and by brain macrophages both positively regulate the formation of CD8+ bTr, with a more pronounced effect on the TP subset. Moreover, the bi-functional effector capacity of CD8+ bTr is enhanced by neuronal MHC I presentation.

**Figure 5.**
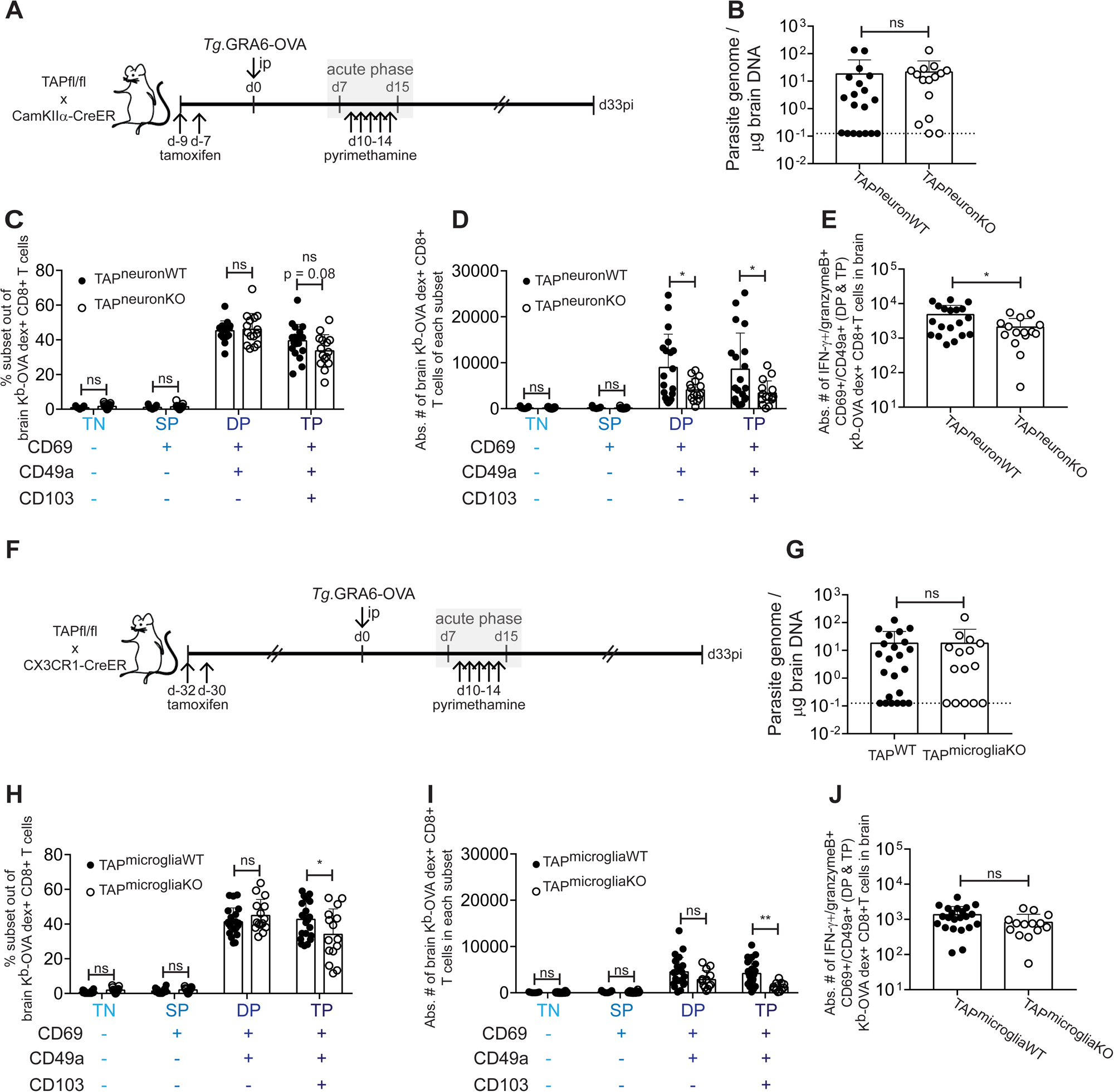
TAP-mediated MHC I antigen presentation by excitatory neurons and CX3CR1+ brain macrophages fine-tune the differentiation of parasite-specific brain-resident CD8+ T cells. (A) Schematics of experimental workflow: Tap1fl/fl X Camk2aCreER+ (TAP^neuronWT^) and Camk2aCreER– (TAP^neuronKO^) mice were treated with tamoxifen, infected one week later with 200 tachyzoites of GRA6-OVA-expressing *T. gondii* Pru, and treated from d10pi to d14pi with an anti-parasitic drug (pyrimethamine) to avoid excessive parasite burden in TAP^neuronKO^ mice due to defective MHCI neuronal presentation (Salvioni et al., 2019). (B) Brain parasite burden measured by qPCR on genomic DNA extracted from brain at d33pi. Mann-Whitney test between TAP^neuronWT^ and TAP^neuronKO^ groups. (C) Graph showing the percentage of each subset (TN: CD69- CD49a- CD103-, SP: CD69+ CD49a- CD103-, DP: CD69+ CD49a+ CD103-, TP: CD69+ CD49a+ CD103+) out of parasite (OVA)-specific CD8+ T cells. Unpaired t-tests performed for each subset between TAP^neuronWT^ and TAP^neuronKO^ groups. (D) Graph showing the absolute number of parasite (OVA)-specific CD8+ bTr cells in each subset. Unpaired t-tests performed for each subset between TAP^neuronWT^ and TAP^neuronKO^ groups. (E) Graph showing the absolute number of parasite (OVA)-specific CD8+ bTr cells (i.e. DP & TP) co-expressing IFN-γ and granzyme B following PMA/ionomycin stimulation, i.e. bi-functional cells. Unpaired t-test performed between TAP^neuronWT^ and TAP^neuronKO^ groups. (B, C, D, E) Outliers were removed with ROUT method with max desired FDR (Q) set at 2% before applying the statistical tests. Bars show mean ± SD of N = 19 *vs*. 15 mice per group, pooled from 3 independent experiments. (F) Schematics of experimental workflow: Tap1fl/fl X Cx3cr1CreER+ (TAP^microgliaWT^) and Cx3cr1CreER– (TAP^microgliaKO^) mice were treated with tamoxifen, infected one month later with 200 tachyzoites of GRA6-OVA-expressing *T. gondii* Pru, and treated from d10pi to d14pi with an anti-parasitic drug (pyrimethamine) to avoid differential parasite burden between the two genotypes. (G) Brain parasite burden measured by qPCR on genomic DNA extracted from brain at d33pi. Bars show the mean of N = 23 *vs*. 15 mice per group, pooled from 3 independent experiments. Mann-Whitney test between TAP^microgliaWT^ and TAP^microgliaKO^ group. (H) Graph showing the percentage of each subset out of parasite (OVA)-specific CD8+ T cells. (I) Graph showing the absolute number of parasite (OVA)-specific CD8+ bTr cells in each subset. (J) Graph showing the absolute number of parasite (OVA)-specific CD8+ bTr cells (i.e. DP & TP) co-expressing IFN-γ and granzyme B following PMA/ionomycin stimulation, i.e. bi-functional cells. (G, H, I, J) Outliers removed with ROUT method with max desired FDR (Q) set at 2%. Bars show mean ± SD of N = 23 *vs*. 15 mice per group, pooled from 3 independent experiments. For all datasets, normality was assessed with D’Agostino & Pearson test. If normal, a two-tailed unpaired t-test was applied, if not, a Mann-Whitney test was chosen.

At last, we decided to take advantage of this latent infection setting to identify subsets of brain-resident CD8+ T cells that may be specifically associated with long-term parasite control and resistance to TE. In order to perform a comparison with a context of susceptibility to TE, we used a Pru-derived *T. gondii* line expressing another OVA-based model antigen (SAG1-OVA) that is more poorly presented than GRA6-OVA and leads to less effective parasite control in the brain (Salvioni et al., 2019; Schaeffer et al., 2009) (**Sup. Fig. 3A**). Whereas mice infected with *Tg*.GRA6-OVA parasites regained weight following acute phase, mice infected with *Tg*.SAG1-OVA parasites developped a more severe form of chronic infection with higher parasite load persisting in the brain following the resolution of acute phase (**Sup. Fig. 3B, 3C**). Using these 2 models, we performed a longitudinal single-cell RNA-seq analysis of parasite (OVA)-specific CD8+ T cells isolated from the brain at different time points post-infection. We Facs-sorted OVA-specific CD8+ T lymphocytes from the brain at ‘early’ (52 days post-infection (dpi)) and ‘late’ (160dpi, ∼5.5 months pi) chronic infection. For ethical reasons, only latently infected mice, and not encephalitic mice, could be included at the late time point. We Facs-sorted parasite-specific CD8+ T cells into CD103-negative and CD103-positive populations (**Sup. Fig. 3D**) and subjected these cells to scRNA-seq processing with the 10x Genomics platform. Following successive steps of quality control, the gene expression profiles of 6182 OVA-specific CD8+ T cells pooled from the 3 conditions (i.e. early encephalitis d52pi, early latency d52pi, late latency d160pi) were integrated with Seurat and included in subsequent analyses (**Sup. Fig. 3E**).

After projection on a Uniform Manifold Approximation and Projection (UMAP) plot, parasite-specific CD8+ T cells partitioned in 13 clusters (**Fig. 6A, 6B**). Six (6) out of 13 clusters (clusters 0, 1, 2, 7, 8, 12 comprising 65 % of the dataset) had a positive enrichment score for at least one published Trm signature and a negative enrichment score for circulating T cell signatures (**Fig. 6C**), confirming that a majority of parasite-specific CD8+ T cells present in the brain during chronic stage indeed have a resident transcriptional profile. Three (3) out of 13 clusters (clusters 3, 5 and 11 comprising 19 % of the dataset) had a positive enrichment score for circulating T cell signatures and a negative enrichment score for Trm signatures, and the 4 remaining clusters (clusters 4, 6, 9, 10 comprising 16 % of the dataset) displayed a mixed profile (**Fig. 6C**). Based on these analyses, we tagged cells within these clusters respectively as resident (bTr), circulating (bTcirc) or mixed (bTmixed).

**Figure 6.**
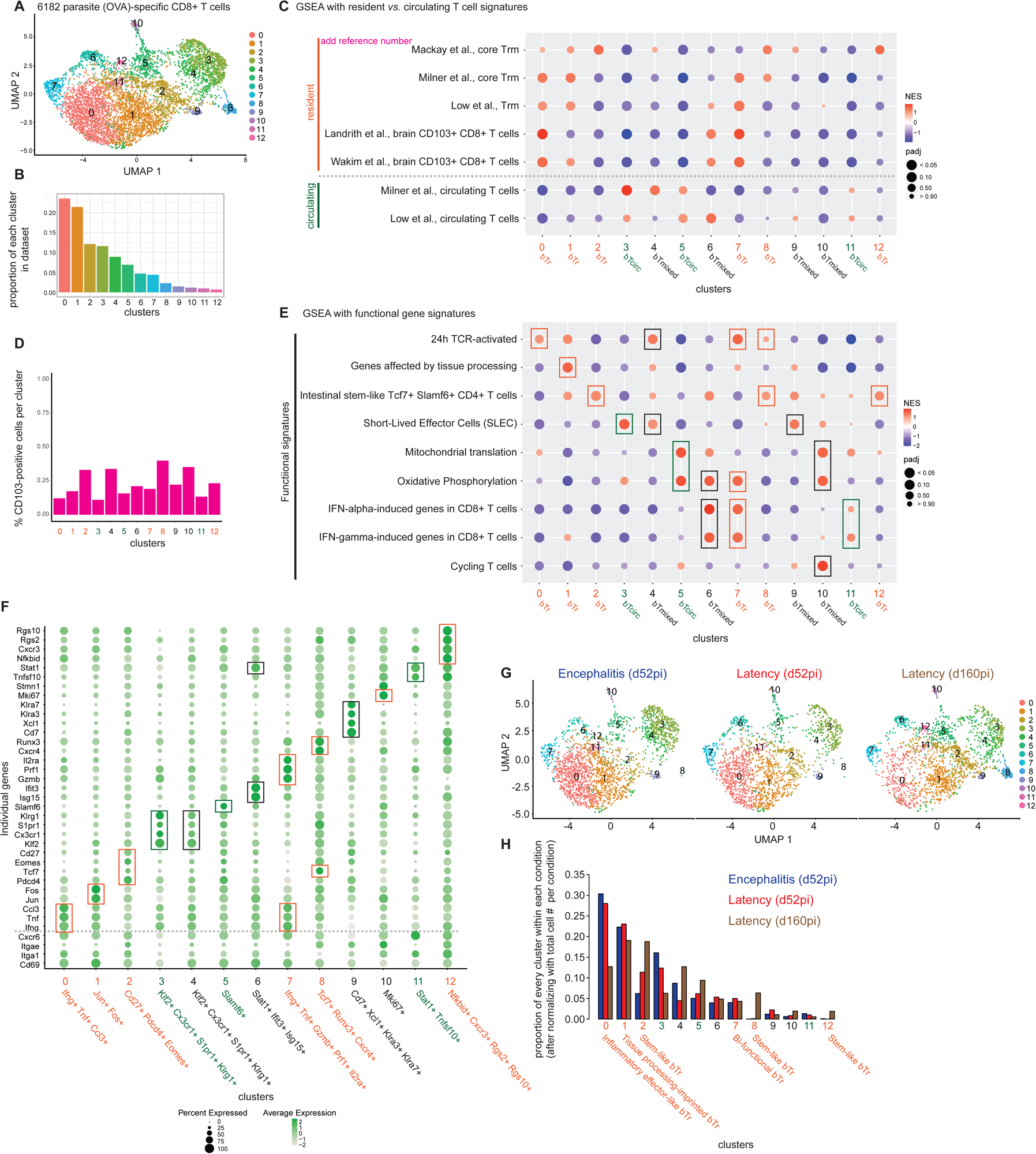
Longitudinal single-cell RNA-seq analysis of brain-isolated *T. gondii*-specific CD8+ T cells in encephalitis and latency infection models. (A) Uniform Manifold Approximation and Projection (UMAP) plot of 6182 brain-isolated OVA-specific CD8+ T cells pooled from 3 conditions: early encephalitis (d52pi, 4 mice pooled), early latency (d52pi, 6 mice pooled) and late latency (d160pi, 10 mice pooled), partitioned in 13 clusters using Seurat-embedded Louvain clustering algorithm. (B) Bar graph showing the proportion of each cluster within entire dataset. (C) GSEA using tissue-resident T cell gene signatures (from (Mackay et al., 2013), (Milner et al., 2017), (Low et al., 2020), (Landrith et al., 2017), (Wakim et al., 2012)) *vs*. circulating T cell gene signatures (from (Milner et al., 2017) and (Low et al., 2020)). Each cluster is colored and tagged as ‘resident’ (bTr, orange) or ‘circulating’ (bTcirc, green), based on the enrichment scores of resident *vs.* circulating T cell signatures. Clusters showing positive enrichment with both types of signatures (hybrid profile) were designated as ‘bTmixed’ (black-colored text). (D) Bar graph showing the proportion of CD103-positive cells (based on Facs-sorting) among each cluster. (E) GSEA using previously published and/or public ‘functional’ gene signatures including: recent TCR activation (Low et al., 2020), imprinting of tissue dissociation procedure (van den Brink et al., 2017), stem-like CD4+ T cells (Schnell et al., 2021), short-lived effector cells (SLEC) (Joshi et al., 2007), mitochondrial translation (Reactome pathway knowledgebase (Gillespie et al., 2022)), oxidative phosphorylation (KEGG pathway database (Kanehisa et al., 2017)), type I and type II IFN responses (Mostafavi et al., 2016), proliferation (Clarke et al., 2019). (F) Dot plot showing average expression (color intensity) and percentage of gene-expressing cells (dot size) per cluster for a panel of individual genes. Boxes around the dots and annotations below the cluster number highlight upregulated genes of interest of every cluster. (G) UMAP separately showing cells from the 3 experimental conditions. (H) Bar graph showing the proportion of each cluster per condition, normalized with respect to the total number of cells analyzed per condition. Annotations below the cluster number indicate the inferred cell ‘identity’.

This first basic GSEA analysis indicated that parasite-specific CD8+ bTr span 6 clusters, and that they are transcriptionally heterogeneous. As expected, the frequency of CD103-positive cells (based on Facs sorting) tended to be lower in bTcirc clusters (#3, #5, #11), but CD103-positive cells did not make up a distinct cluster of bTr (**Fig. 6D**). CD103-positive cells partitioned in bTr and non-bTr clusters in variable proportions, emphasizing the notion that CD103 is not a universal marker of tissue-resident T cells (Bergsbaken and Bevan, 2015; Bergsbaken et al., 2017; Ferreira et al., 2020; Steinert et al., 2015). Additional GSEA for ‘functional’ gene signatures (**Fig. 6E**) and inspection of individual marker genes (**Fig. 6F**) further underlined the marked diversity of *T. gondii*-specific CD8+ bTr and allowed to infer their functional status. We identified (i) “inflammatory effector” bTr cells (cluster 0) that have recently been activated by the TCR and overexpress Tnf, Ifng and chemokine genes, (ii) “stem-like” bTr cells (clusters 2, 8 & 12), which display lower expression of cytokine genes and higher expression of genes associated with stemness in T cells (e.g. Cd27, Tcf7), which are transcriptionally close to progenitor CD4+ T cells (Schnell et al., 2021) and which, especially for cluster 2, overexpress Pdcd4, a translational repressor involved in the downregulation of IFN-γ production in T cells (Lingel et al., 2017), (iii) “bi-functional” bTr cells (cluster 7), which have also been recently TCR-activated, which display IFN-responsive signatures, and which simultaneously overexpress Ifng and cytotoxicity-associated genes (e.g. Prf1 & Gzmb), indicative of a dual effector activity and a potential implication in parasite containment. In addition, some bTr cells (grouped within cluster 1) showed upregulation of Jun and Fos early response genes, as well as of many other genes known to be affected by the cell isolation procedure (Crowl et al., 2022; van den Brink et al., 2017). We called cells within this cluster “tissue-processing imprinted”. Remarkably, none of the 13 clusters upregulated the typical T cell exhaustion genes (e.g. Pdcd1 and Tox), confirming the absence of T cell functional exhaustion, a feature that was previously reported in in another model of latent *T. gondii* infection using adoptively transferred parasite-specific CD8+ T cells (Chu et al., 2016). By examining the temporal evolution of these 13 clusters (**Fig. 6G, 6H**), we noticed that the abundance of “bi-functional” bTr cells (cluster 7) and of “tissue processing-imprinted” cells (cluster 1) was largely similar across the conditions. Expectedly, we found that “inflammatory effector” bTr cells (cluster 0) predominated during encephalitis and early latency, and decreased upon late latency. In contrast, “stem-like” bTr cells (clusters 2, 8 & 12) inflated over time and were most abundant at late latency. To confirm the existence and temporal evolution of such stem-like subsets, we used flow cytometry to monitor the proportion of TCF1 (Tcf7 gene)/CD27 double-positive cells among CD69+ CD49a+ CD8+ bTr throughout early (d37pi) and late (d135pi) latency (**Sup. Fig. 4A, B, C**). This complementary read-out confirmed that stem-like T cells indeed make up an increasing proportion of bTr, both among parasite (OVA)-specific CD8+ T cells (**Sup. Fig. 4D**) and among total CD8+ T cells (**Sup. Fig. 4E**), suggesting that they might be important contributors of long-term parasite control during latency. The above data were obtained with endogenous CD8+ T cells specific for OVA-based model antigens expressed by *T. gondii*. To confirm that CD8+ T cells specific for a natural parasite antigen display a similar transcriptional diversity, we infected H-2L^d^-transgenic C57BL/6 mice (B6.L^d^ (Salvioni et al., 2019)) with Tg.GRA6-OVA and performed a new single-cell transcriptomic analysis of brain-isolated CD8+ T cells at early latency (46 dpi) (**Sup. Fig. 5A**). In addition to OVA-specific CD8+ T cells, infection of this B6.L^d^ transgenic mouse model with type II *T. gondii* elicits CD8+ T cells that recognize a decameric antigenic peptide (HF10) that is naturally processed from the GRA6 protein of *T. gondii* and is presented by H-2L^d^ MHC I (Blanchard et al., 2008; Salvioni et al., 2019). The analysis of 2715 total CD8+ T cells isolated from the brain revealed 9 clusters (**Sup. Fig. 5B, 5C**). GSEA with the same resident *vs.* circulating T cell gene signatures as above (see Fig. 6) showed that half of such CD8+ T cells (∼49%, corresponding to clusters 0, 1, 3 & 8) could be defined as bTr based on their preferential enrichment in Trm signatures, a quarter of the cells (∼24%, corresponding to clusters 2 & 5) could be defined as Tcirc based on their enrichment in circulating T cell signatures, and the rest (∼27%, clusters 4, 6, 7) had unclear and/or hybrid transcriptional profiles, and were tagged as bTmixed (**Sup. Fig. 5D**). Thanks to barcoded (dCode) MHC class I dextramers loaded with the OVA-derived (K^b^-OVA) or GRA6-derived (L^d^-GRA6) peptides, we used CITE-seq to retrospectively identify CD8+ T cells specific for OVA or GRA6 in each cluster (**Sup. Fig. 5E, F**). Based on the overall abundance of L^d^-GRA6- vs. K^b^-OVA-labelled T cells, the GRA6-specific response appeared subdominant compared to the OVA-specific response. Nevertheless, T cells of both antigenic specificities distributed in all clusters, with the exception of cluster 7, which corresponds to Pdcd1+ Tox+ exhausted T cells (**Sup. Fig. 5G**). This indicated that T cell exhaustion does occur during latent *T. gondii* infection but not in the parasite-specific compartment. As above, we plotted the average expression of the same set of individual genes (**Sup. Fig. 5G**) and performed GSEA with the same functional gene signatures on these 9 clusters (**Sup. Fig. 5H**). To help highlight the equivalence between clusters identified in this new experiment and clusters from the former experiment performed with OVA-specific CD8+ T cells (depicted in Fig. 6), we calculated the Spearman correlation coefficient between clusters of each experiment, based on the log2 fold-change of all genes expressed by cells within each cluster (**Sup. Fig. 5I**). This set of analyses allowed us to identify one subset of tissue processing-imprinted bTr cells (cluster 3), one subset of Ifng+ Gzmb+ bi-functional bTr (cluster 8), one subset of TCR-activated Runx3+ Pdcd4+ bTr (cluster 0), and two clusters of “stem-like” CD8+ T cells (one of bTr: cluster 1, and one of bTmixed: cluster 4).

In conclusion, single-cell transcriptomic analyses of CD8+ T cells isolated from the brain confirmed that a majority of parasite-specific CD8+ T cells display tissue-resident transcriptional profiles and that they are refractory to functional exhaustion. In addition, these analyses unveiled the high transcriptional diversity of parasite-specific CD8+ bTr cells, ranging from progenitor-like to bi-functional effector phenotypes. They showed that long-term parasite control in the brain (i.e. late latency) is associated with the expansion of TCF1+ CD27+ CD8+ bTr that display stem-like features.

## Discussion

Through a combination of single-cell approaches, antibody-based T cell depletion strategies, and a novel conditional KO model to cripple the MHC I antigen processing pathway in brain antigen-presenting cells (APC), our work sheds light on 3 major unresolved questions during *T. gondii* latent infection: 1) the role of brain-resident CD8+ T cells in parasite immune surveillance, 2) the cellular cues that govern the differentiation of such brain-resident CD8+ T cells, and 3) their functional diversity.

A key achievement of our work is to address how MHC I presentation of parasite antigens by excitatory neurons and resident macrophages within the brain regulates the formation of CD8+ bTr. While trafficking of activated CD8+ T cells into a tissue is mostly inflammation-driven, the differentiation into Trm is thought to be mostly controlled by local cues. Studies have shown that cognate antigen recognition within the tissue is a major driver of Trm differentiation and survival. For example, antigen stimulation in the skin upon Vaccinia virus infection induces CD69 and promotes skin retention of cognate CD8+ T cells (Khan et al., 2016). Antigen stimulation in the brain upon VSV infection was reported to promote CD103 expression on CD8+ bTrm (Wakim et al., 2010). Regarding the respective contributions of tissue APC, a pioneer study in the lung, in the context of chronic LCMV infection, showed that although leading to different transcriptional responses, both hematopoietic and non-hematopoietic APC of the lung were able to reactivate lung CD8+ Trm upon re-exposure to the virus (Low et al., 2020). This was in contrast to the lung-draining lymph nodes where memory CD8+ T cells were strictly reliant on DC for their reactivation (Low et al., 2020). To our knowledge, conditional KO approaches to genetically dissect the respective contributions of tissue APC in resident T cell differentiation have never been used so far. The model implemented for this study allowed cell type-specific conditional invalidation of TAP1, a critical component of MHC I antigen processing, known to be important for presentation of the GRA6-derived antigenic peptide (Blanchard et al., 2008). Neurons are the main target cells of *T. gondii* in the brain (Cabral et al., 2016) and the only reservoir of parasite cysts during chronic stage (Melzer et al., 2010). Moreover, our previous work showed that MHC I presentation of tachyzoite-derived antigens by excitatory neurons during early chronic stage (∼d30pi) drives resistance against TE, without affecting the accumulation of CD8+ T cells in the CNS (Salvioni et al., 2019). Beside neurons, microglia are also relevant host cells for *T. gondii* (Cabral et al., 2016) and their ability to respond to IFN-γ is instrumental for optimal parasite control in the brain (Cowan et al., 2022). Based on this, we decided to investigate the role of both types of APC in the formation of CD8+ bTrm using tamoxifen-inducible Cre lines. Overall, our data show that MHC I presentation by neurons and microglia has modest but clear effects on the differentiation of parasite-specific CD8+ bTr. In line with a former study of brain CD8+ Trm upon viral infection showing a requirement for antigen recognition within the brain for expression of CD103 on Trm (Wakim et al., 2010), we observed that TAP-dependent MHC I presentation of *T. gondii*-derived antigens by both excitatory neurons and microglia promotes the generation of TP (CD69+ CD49a+ CD103+) CD8+ bTr. An additional and intriguing result is that the generation of bi-functional CD8+ bTr relied preferentially on the ability of T cells to recognize their cognate antigen on the surface of excitatory neurons. These differential requirements may be linked to the quantity of peptide-MHC complexes present on the surface of each APC type or, perhaps more interestingly, on the parasite stage from which the antigen is processed (assuming that only neurons could process and present bradyzoite-secreted GRA6-OVA). Whether, and to which extent, CD8+ bTrm differentiation is shaped by parasite effectors that may manipulate host responses in a stage-specific manner, remains to be investigated.

A second critical finding of our work is to uncover the protective function of *T. gondii*-specific CD8+ bTr. While CD8+ T cells with a phenotypic and transcriptional profile reminiscent of Trm had been previously reported in the brain of mice chronically infected with *T. gondii* (Landrith et al., 2017; Sanecka et al., 2018), the function of these resident T cells had remained unstudied. Our C57BL/6-based infection model mimicking latent infection provided a well-suited context to examine the functional role of CD8+ bTr. Up to now, no genetic model allowing a targeted elimination of brain-resident T cells has been published. Our attempts to use the Hobit-tdTomato-DTR model to eliminate Hobit+ T cells via diphtheria toxin (DT) injection failed for brain-resident cells. As described previously (Parga-Vidal et al., 2021; Zundler et al., 2019), we confirmed that DT injection reduces Trm numbers in the liver and the intestine, but this treatment was ineffective in the brain (not shown).

To overcome this limitation, we had to resort to using indirect approaches. Injection of anti-CD8 antibodies during chronic infection, which leaves intact the tissue-resident T cell compartment but cripples the circulating CD8 T cells (Frieser et al., 2022; Steinbach et al., 2016), showed that circulating CD8+ T cells are dispensable for optimal parasite control in the brain while they limit the recrudescence of *T. gondii* in the spleen. Based on our current understanding of latent *T. gondii* infection, the only reservoir of parasites during latency is the CNS. Hence, we assume that the increased parasite load observed in the spleen upon CD8 T cell depletion is not linked to a local (splenic) reactivation but rather due to the continuous escape of a few parasites (most likely tachyzoïtes) from the CNS to the periphery, possibly through meningeal lymphatics (Kovacs et al., 2022). We hypothesize that in the absence of peripheral CD8+ T cells, these ‘escaped’ tachyzoites were poorly controlled and thus able to replicate, leading to their detectability by qPCR two weeks post-depletion. The protective role of bTr was revealed as well when we depleted the CD4 T cell compartment. The fact that, in mice, CD4+ T cells play a limited role in parasite restriction and are not mandatory to survive acute stage (Schaeffer et al., 2009), allowed us to focus on the impact of CD4+ T cells during chronic stage. In CD4+ T cell-depleted mice, we observed a strong blockade in the differentiation of CD8+ bTr from the CD69+ CD49a+ double-positive (DP) into the CD69+ CD49a+ CD103+ triple-positive (TP) cells, correlating with an increase in brain parasite burden. This result suggests that the CD4+ T cell-dependent subset of TP CD8+ bTr strongly contributes to keeping the parasites in check in the brain. Noteworthy, our experimental set-up (in which CD4+ T cells are depleted prior to infection) did not allow us to conclude if the CD4+ T cell-mediated parasite restriction is set early at the time of parasite invasion and is durable, or if CD4+ T cell help is constantly needed for optimal maintenance of the resident CD8+ T cell compartment. Regardless, our data emphasizing the critical role of CD4+ T cells in maintaining effector function of CD8+ T cells and in *T. gondii* control, appear in agreement and extend previous findings from the Khan laboratory, who reported the importance of CD4+ T cells in maintaining the functionality of CD8+ T cells and showed in a pioneer study that the IL-21 receptor is required for the control of *T. gondii* reactivation in the brain (Moretto et al., 2017). At the time of their publication, the concept of resident T cells was just emerging. Their findings might now be reinterpreted in the light of the importance of IL-21 for tissue-resident CD8+ T cell differentiation (Ren et al., 2020). Importantly, we think that our data help understand the long known association between HIV/AIDS and parasite reactivation leading to TE in humans (Nissapatorn, 2009). During HIV infection, HIV is known to infect and elicit death of CD4+, not CD8+, T cells. Nevertheless, both the CD4+ and CD8+ T cell compartments display major dysfunctions upon HIV infection (Fenwick et al., 2019). Such dysfunctions include not only the well-studied state of functional exhaustion (Day et al., 2006) but also an impairment in Trm differentiation and maintenance. Accordingly, it was reported that HIV-infected individuals presenting with a low CD4+ T cell nadir (i.e. that have started anti-retroviral therapy at a late timepoint) display an irreversible reduction and dysregulation of mucosal tissue-resident CD4+ and CD8+ T cells, leading to weakened immune control of human papilloma virus (HPV) in the skin and increased development of HPV-related cancer (Saluzzo et al., 2021). It is tempting to speculate that alterations of the CD4+ T cell compartment during HIV/AIDS may also impair resident memory CD8+ T cells in the CNS, leading to subpar *T. gondii* immune surveillance in the brain and TE development. Future studies examining resident T cells in brain biopsies of TE patients could be helpful to address this question.

A third asset of our study is to solve an apparent paradox about T cell exhaustion during chronic *T. gondii* infection. While T cell exhaustion has been observed in both CD4+ (Hwang et al., 2016) and CD8+ T cells (Bhadra et al., 2011) during chronic infection in TE models, in a latent infection model, a robust effector CD8+ T cell response was shown to be maintained over time without signs of functional exhaustion (Chu et al., 2016). By tracking CD8+ T cells specific for 2 distinct parasite-derived antigenic peptides in single-cell RNA-seq clusters, our CITE-seq analysis revealed that parasite-specific CD8+ T cells are absent from the cluster containing exhausted CD8+ T cells (see cluster 7 on Sup. Fig. 5). This indicated that functional exhaustion of CD8+ T cells does indeed occur in the brain during latent infection, but not in the *T. gondii*-specific CD8+ T cells. It is likely that this unexpected and original phenomenon had been overlooked so far, since the Khan study (Bhadra et al., 2011) did not distinguish between bystander and parasite-specific CD8+ T cells. Understanding the mechanisms that shield parasite-specific CD8+ T cells from exhaustion could offer new therapeutic options to improve T cell functionality in contexts of tissue-restricted antigen persistence such as chronic infections or cancer (Baessler and Vignali, 2024; Hashimoto et al., 2023).

Importantly, we provide evidence that parasite-specific CD8+ T cells stationed in the brain play an essential and largely autonomous role in the restriction of *T. gondii* in the brain. This concept of autonomous cellular compartment is in agreement with recent work showing that meningeal lymphatic drainage from the brain supports the development of peripheral T cell responses against *T. gondii* but is dispensable for immune protection of the brain (Kovacs et al., 2022). Notably however, our work did not evaluate the importance of *T. gondii*-induced Trm in other tissues. Interrogating how parasite-specific CD8+ Trm are formed in the small intestine, which is the typical entry site of the parasite, and studying their implication as a first line of defense upon a new encounter with cysts or oocysts, represent exciting avenues of future investigations.

At last, our study contributed to unveil the marked functional diversity of CD8+ bTr. We identified a subset of cytokine/chemokine-producing cells that were recently activated by the TCR. This subset represents the most prominent cluster at d52pi in the encephalitis and latency contexts, and it subsides with time in the late latency condition. Cells within this cluster likely overlap with *T. gondii*-specific Nur77-GFP+ CD8+ T cells undergoing secondary TCR engagement, which were described in a recent study of the Hunter laboratory following adoptive transfer of TCR-transgenic OT-I CD8+ T cells, and were observed in association with areas of tachyzoite replication in the brain (Shallberg et al., 2022). Interestingly, among the bTr clusters that underwent recent TCR activation (see clusters 0, 7, 8 on Fig. 6), we found a “bi-functional” subset simultaneously overexpressing Ifng and cytotoxicity-associated genes (e.g. Prf1 & Gzmb), leading us to suspect that these cells might play a predominant role in parasite control. Testing their functional role will require the set-up of selective methods to target resident subpopulations in the brain. An exciting finding of our transcriptomic analyses is the discovery of bTr subsets which expand over time in the brain of latently infected mice and ultimately represent more than one-third of the parasite-specific CD8+ T cells at 5.5 months pi (see clusters 2, 8, 12 on Fig. 6 and Sup. Fig. 4). These cells are transcriptionally close to stem-like (Tcf7+ Slamf6+) CD4+ T cells that were found in the intestine and serve as a reservoir of extra-intestinal effector CD4+ T cells with an encephalitogenic potential (Schnell et al., 2021). Based on the expression of genes associated with T cell stemness, such as the transcription factor Tcf7, we hypothesize that such progenitor CD8+ bTr may display an intrinsic self-renewal capacity and contribute to durably fuel the pool of more effector brain-resident CD8+ T cells that are needed to circumvent parasite reactivation. Interestingly, in a context of chronic systemic viral infection (LCMV), IL-33 signals have been shown to control the expression of Tcf7, thereby promoting the expansion of stem-like peripheral CD8+ T cells (Marx et al., 2023). Since IL-33 is released by oligodendrocytes and astrocytes in the brain during *T. gondii* infection (Still et al., 2020), a similar pathway may be at play in the maintenance of brain-resident progenitor T cell subsets. Adequate tools will need to be developped to specifically interrogate the role of IL-33 signaling in the generation/maintenance of stem-like CD8+ bTr, and to determine if these cells could be a self-renewing reservoir of bTr. Altogether, our findings suggest that any strategy aimed at boosting the CD8+ Trm compartment, could be useful to mitigate the risk of parasite reactivation in immunocompromised individuals.

## Materials & Methods

### Mice

Animal care and use protocols were carried out under the control of the French National Veterinary Services and in accordance with the current European regulations (including EU Council Directive, 2010/63/EU, September 2010). The protocol “APAFIS#25130-2020040721346790 v3” was approved by the local Ethical Committee for Animal Experimentation registered by the “Comité National de Réflexion Ethique sur l’Experimentation Animale” under no. CEEA122. C57BL/6J (B6) mice were purchased from Janvier (France). H-2 L^d^-transgenic C57BL/6J (B6.L^d^), Hobit^tdTomato-DTR^ (Parga-Vidal et al., 2021), Camk2a-CreER (Casanova et al., 2001) and Cx3cr1-CreER (Parkhurst et al., 2013) mice were previously described. Tap1^fl/fl^ mice were obtained by crossing conditional-ready C57BL/6N-Tap1^tm2a(EUCOMM)Hmgu^/Ieg mice generated by the European Conditional Mouse Mutagenesis Program(EUCOMM EM:09400 strain) with FlpO-deleter mice (C57BL/6-Tg(CAG-Flpo)1Afst/Ieg (Kranz et al., 2010), EUCOMM EM:05149 strain) in order to achieve flippase-mediated excision of the LacZ and Neomycine resistance cassette. All non-commercial mouse models were housed and bred under specific pathogen-free conditions at the ‘Centre Regional d’Exploration Fonctionnelle et de Ressources Experimentales’ (CREFRE-Inserm UMS006). Mice were experimentally infected between 8 and 12 weeks of age. All mice used in experiments were males. Number of mice and experimental replicates are indicated in the respective figure legends.

### Human cell lines

Male human Foreskin Fibroblasts (HFF) were purchased from ATCC (Hs27 ref # CRL-1634). HFF were maintained in DMEM supplemented with 10% FCS (GIBCO ref # 10270106).

### Toxoplasma gondii

Mouse infections were done intra-peritoneally (i.p.) with tachyzoites of GFP+ type II Prugnaud (Pru) *T. gondii*, expressing GRA6-OVA under the control of the endogenous GRA6 promoter (Pru.GRA6-OVA (Salvioni et al., 2019)) or tachyzoites of tdTomato+ Pru *T. gondii* expressing SAG1-OVA (Pru.SAG1-OVA (Schaeffer et al., 2009)). Tachyzoites were maintained *in vitro* by serial passages on confluent monolayers of HFF using DMEM supplemented with 1% FCS (GIBCO). For mouse infections, infected HFF were scraped, tachyzoites were released through a 23G needle, filtered through a 3 µm polycarbonate hydrophilic filter (it4ip S.A.) and 200 tachyzoites were injected i.p. in 200µL PBS.

### Antibody-based T cell depletion

To eliminate circulating CD8+ T cells during chronic phase, a depleting anti-CD8β mAb (rat IgG1, Lyt3.2, clone 53-5.8, cat # BX-BE0223 from BioXcell) was injected i.p. for 2 weeks, starting at 34 and 36 days post-infection (dpi) (200 μg per mouse), and then once at d41pi and once at d48pi (100 μg per mouse), before euthanasia at d50pi. To deplete CD4+ T cells, a depleting anti-CD4 mAb (rat IgG2b, clone GK1.5, cat # BX-BP0003 from BioXcell) was injected i.p. at day 3 (200 μg per mouse) and day 1 (100 μg per mouse) prior to infection, and then once weekly (100 μg per mouse) until euthanasia at d13pi (acute phase analysis) or d33pi (chronic phase analysis). Control mice were injected with the same amount of isotype control rat antibodies (rat IgG1, clone HRPN, cat # BX-BE0088 from BioXCell and rat IgG2b, clone LTF-2, cat # BX-BP0090 from BioXCell respectively).

### Tamoxifen and pyrimethamine treatments

Tamoxifen (Sigma-Aldrich, cat # T5648) was dissolved in corn oil at a concentration of 50 mg/mL, and 200 µl were administered twice *per os* two days apart (i.e. 9 and 7 days prior to infection of Tap1^fl/fl^ x CamK2a-CreER mice, or 32 and 30 days prior to infection of Tap1^fl/fl^ x Cx3cr1-CreER mice). A 10X pyrimethamine (Sigma-Aldrich cat # 46706) solution (12,5 mg/mL) was prepared in DMSO (Sigma-Aldrich cat # D2650) and stored at 4°C until diluted 10-fold in corn oil just before administration. A dose of 12,5 mg/kg/day of pyrimethamine was administered daily *per os* for 5 days, starting at d10pi.

### Isolation of spleen and brain leukocytes

Spleens and brains were collected in complete RPMI (GIBCO) supplemented with 10% vol/vol FCS (GIBCO). Spleens were mashed through a 100 μm cell strainer (Falcon). Brains were homogenized using a glass Potter and digested for 45 min at room temperature (RT) in Hanks’ balanced salt solution (HBSS) medium with collagenase D (1 mg/ml, Roche Diagnostics cat # 11088882001) and deoxyribonuclease (DNase) I (20 μg/ml, Sigma-Aldrich cat # DN25). After digestion, cells were suspended in 30% Percoll (GE Healthcare) and centrifuged at 1590g for 30 min without brake. Myelin and debris were removed, brain leukocytes were recovered from the interface and further used for experiments. Erythrocytes were lyzed from brain and spleen cell suspensions using ACK buffer (100 mM EDTA, 160 mM NH4Cl and 10 mM NaHCO3).

### *Ex vivo* T cell restimulation

Brain or spleen cells were plated in a V-bottom 96 well plate and cultured in complete RPMI supplemented with 0.05 µg/mL Phorbol 12-myristate 13-acetate (PMA), 1 µg/mL ionomycine and 3 µg/mL brefeldin A for 4h at 37°C, 5% CO_2_. Cells were then washed once in complete RPMI before performing flow cytometry stainings.

### Antibody stainings for flow cytometry and/or cell sorting

To detect CD8+ T cells specific for the OVA-derived SIINFEKL peptide presented by H-2K^b^, splenocytes and brain leukocytes were incubated for 45 min at RT with PE-coupled SIINFEKL-loaded H-2 K^b^ dextramers (Immudex, dilution 1:50 in complete RPMI).

For flow cytometry, following Fc receptor saturation (CD16/32, clone 93, dilution 1:50 in PBS, Biolegend) and dead cell detection with Fixable Viability Stain 440UV (dilution 1:500 in PBS, BD Horizon cat # 566332), cell suspensions were surface labelled for 30 min at 4°C with CD3 BV785 (clone 17A2, 1:50, BD Horizon), CD8α BUV805 (clone 53-6.7, 1:800, BD Horizon), CD4 APC-Cyanine7 (clone GK1.5, dilution 1:100, BD Pharmingen) or CD4-AF700 [clone RM4-5, dilution 1:200, BD Pharmingen], CD49a BUV737 (clone Ha31/8, dilution 1:400, BD OptiBuild), CD69 BUV615 (clone H1.2F3, dilution 1:50, BD OptiBuild), CD103 BV510 (clone M290, dilution 1:200, BD Horizon), CD27 BV421 (clone LG.3A10, dilution 1:200, BD Optibuild ref 740028). Fluorochrome-coupled antibodies were diluted in Brilliant Stain Buffer (BD Horizon). Intracellular IFN-γ BV421 (clone XMG1.2, 1:100, BD Horizon), Granzyme B PE-Cyanine7 (clone NGZB, 1:200, eBioscience) and TCF1 AF488 (clone C63D9, dilution 1:200, Ozyme) stainings were performed with Foxp3/Transcription Factor Staining Buffer Set (eBioscience) following the manufacturer’s protocol.

Before scRNA-seq, CD103-negative and CD103-positive subpopulations of K^b^-OVA dex+ CD8+ T cells were separated by Facs-sorting. For CITE-seq, brain cell suspensions were incubated for 45 min at RT with the following barcoded dextramers diluted in complete RPMI: K^b^-SIINFEKL (Immudex cat # JD2163-PfBC, barcode fBC0074: CGGTCTTAGTCGCGC, dilution 1:50), L^d^-HPGSVNEFDF (Immudex cat # JG5820-PfBC0301, barcode fBC0301: CGGCCTCGCGACGAC, dilution 1:50), control K^b^-SIYRYYGL (Immudex cat # JD2164-PfBC, barcode fBC0068: TTGCGCGGCGTCGTA, dilution 1:50), and then surface labelled for 30 min at 4°C with TotalSeq-C0182 anti-mouse CD3 (clone 17A2, dilution 1:50, Biolegend, cat # 100263) and TotalSeq-C0002 anti-mouse CD8α (clone 53-6.7, dilution 1:50, Biolegend, cat # 100785) diluted in ‘Facs buffer’ (PBS, 2mM EDTA, 0.5% FCS).

### Parasite load measurements

For cyst enumeration, 5 % of Potter-dissociated whole brain homogenate was labelled with rhodamine-conjugated Dolichos Biflorus Agglutinin (Vector Laboratories RL-1032). Cysts were counted using an inverted fluorescence microscope with a 20X objective. Quantification of parasite DNA by qPCR was performed on genomic DNA extracted with DNEasy Blood & Tissue Kit (Qiagen) from 5% of each brain homogenate and spleen cell preparation. As described earlier (Feliu et al., 2013), a 529-bp repeat element in the *T. gondii* genome was amplified using the TOX9 and TOX11 primers. The number of parasite genome per μg of tissue DNA was estimated by comparison with a standard curve, established with a known number of Pru tachyzoites. The limit of quantification corresponds to the highest dilution of the standard curve, above which concentrations can be reliably extrapolated.

## Single-cell RNA-sequencing and CITE-seq

### Library preparations for 3’ single-cell RNA-sequencing

Facs-sorted CD103-negative *vs.* CD103-positive CD8+ K^b^-OVA dex+ T cells from the different biological conditions were obtained in two independent experiments. One experiment comprised an encephalitis (Tg.SAG1-OVA-infected C57BL/6) and latency (Tg.GRA6-OVA-infected C57BL/6) group analyzed at d52pi, and one comprised only the latency group analyzed at d160pi. To improve the specificity of the sorting, the antibody panel contained the 3 following markers for a ‘dump’ gate: NK1.1 CD19 MHCII. Single-cell libraries were generated immediately after Facs-sorting using the Chromium Controller Instrument and Chromium Single Cell 3’ Library & Gel Bead Kit v3 according to the manufacturer’s protocol (10X Genomics).

### Library preparations for 5’ single-cell RNA-sequencing (CITE-seq)

CITE-seq dataset is derived from one experiment in which B6.L^d^ mice were infected with Tg.GRA6-OVA and analyzed at d46pi. Brain-isolated CD8^+^ T cells were pooled from 4 mice and labeled with TotalSeqC CD3 and CD8α antibodies (Biolegend), and with PE-coupled dCODE K^b^-OVA, L^d^-GRA6 and Kb-SIYRYYGL control dextramers, to enable retrospective identification of OVA-specific and GRA6-specific cells among total CD8+ T cells. dCODE dextramer-positive CD8+ T cells were enriched by 2 successive magnetic sorting steps: one in which CD8+ T cells were enriched by negative sorting (using Miltenyi Biotec 130-104-075 CD8α+ T cell isolation kit) and one in which PE (dex)+ cells were enriched by positive sorting, using anti-PE microbeads (Miltenyi 130-048-801). Single-cell libraries were generated immediately with Chromium Next GEM Single Cell 5’ Kit v2 and 5’ Feature Barcode Kit according to the manufacturer’s protocol (10X Genomics).

For both 3’ and 5’ libraries, library size and quality were confirmed on a Fragment Analyzer (Agilent). Libraries were sequenced on SP flowcell of Illumina NovaSeq in paired-end sequencing (2 x 150) and a single 8 bp-long index.

### Quality control, dimensionality reduction, and clustering

Cell Ranger software (10X Genomics) (Zheng et al., 2017) was used to perform alignments against *Mus musculus* genome (GRCm38.98 for scRNA-seq and GRCm39.105 for CITE-seq), generating gene expression matrices that were further analyzed with Seurat v4.3.0 (Hao et al., 2021) on R v4.2.2 (R Core Team. R Foundation for Statistical Computing, 2022). A series of filters were applied to remove doublets and low-quality or dying cells. Sample-specific filters were applied on number of genes (nFeature), total number of UMI detected (nCount) and percent counts derived from mitochondrial genes (percent.mt) within a cell. Cell clusters identified by clustifyr package (v1.10.0) (Fu et al., 2020) as ‘non CD8+ T cells’ (e.g. regulatory T cells, macrophages or erythroblasts), as well as the marker genes of these ‘contaminating’ clusters, were omitted from the final dataset. The final CITE-seq dataset was obtained following supplementary filtration steps based on ADT expression, whereby cells expressing aberrantly low levels of CD3 and CD8α, cells positive for the control dCODE dextramer loaded with the irrelevant SIYRYYGL peptide, and cells double positive for K^b^-OVA and L^d^-GRA6 dextramers, were excluded.

Then, Seurat’s integration pipeline for scRNA-seq dataset, and Seurat’s SCTransform function were chosen for normalizing the CITE-seq counts. Variable features were identified and selected (VariableFeatures or FindVariableFeatures and SelectIntegrationFeatures for integration), following by Seurat’s integration functions for scRNA-seq. RunPCA was used to perform linear dimensional reduction based on statistically significant principal components. FindNeighbors, FindClusters, RunUMAP, and FindAllMarkers, FindMarkers functions were executed to infer and visualize cell clusters, and look for differentially expressed genes.

///Briefly, we choose Seurat’s integration pipeline for scRNAseq dataset and SCTranform Seurat’s function for normalize CITEseq counts. RunPCA was used to perform linear dimensional reduction based on statistically significant principal components, and FindNeighbors, FindClusters, RunUMAP, and FindAllMarkers, FindMarkers functions were executed to infer and visualize cell clusters, and look for differentially expressed genes. Functional enrichment analyses were performed with fgsea package v1.24.0 using previously published (see list in figure legends) or publicly available ‘functional’ gene signatures. Spearman’s correlation matrix between clusters was constructed based on log2 fold-change for each gene by cluster, using the correlation computation from the Hmisc package (function rcorr) v5.1-1.

### Statistical analyses

Normality of all datasets was assessed with D’Agostino & Pearson test. If normal, a two-tailed unpaired t-test was applied, if not, a Mann-Whitney test was chosen. Asterisks on graphs reflect statistical significance according to the following standard intervals: **** p < 0.0001, *** p < 0.001, ** p < 0.01, * p < 0.05.

## ACKNOWLEDGEMENTS

We thank R. Balouzat, R. Ecalard, F. Chaboud, E. Debon, M.A. El Manfaloti, S. Negroni, J. Leblond, S. Fresse, M. Lulka from ANEXPLO-CREFRE UMS006 for ethical care of our models, F. Martins & E. Lhuillier from Genotoul GeT-Santé for expert assistance on scRNA-seq, S. Allart, S. Lachambre, L. Lobjois, F. L’Faqihi-Olive, V. Duplan-Eche, A.-L. Iscache, H. Garnier from the microscopy and flow cytometry core facilities of Infinity for technical help. This work was supported by institutional grants from Inserm, PIA PARAFRAP Consortium (ANR-11-LABX0024 to NB), PIA ANINFIMIP equipment (ANR-11-EQPX-0003 to NB), “Agence Nationale pour la Recherche” (ANR-18-CE15-0015 MICCHROB to NB; ANR-19-CE15-0008 TRANSMIT to FM and NB; ANR-22-CE14-0053 NINTENDO to NB), “Fondation pour la Recherche sur le Cerveau” AAP2021 to NB, ANRS0366 to NB. MB was supported by “Fondation Vaincre Alzheimer (FVA)”.

## Contributions

RP, MB, AA, FM, NB designed experiments. RP, MB, AmA, EB, AlA, MA, RMC, AJ performed experiments. RP, MB, AlA, EB, AmA, MA, FM, NB analyzed experiments. KvG generated the Hobit-tdTomato mouse model and provided expert advice. MA & NB carried out bioinformatic analyses. NB wrote the manuscript. All authors read, edited, and approved the final manuscript.

## ETHICAL DECLARATIONS

The authors declare no competing interests.

---

**Sup. Figure 1.**
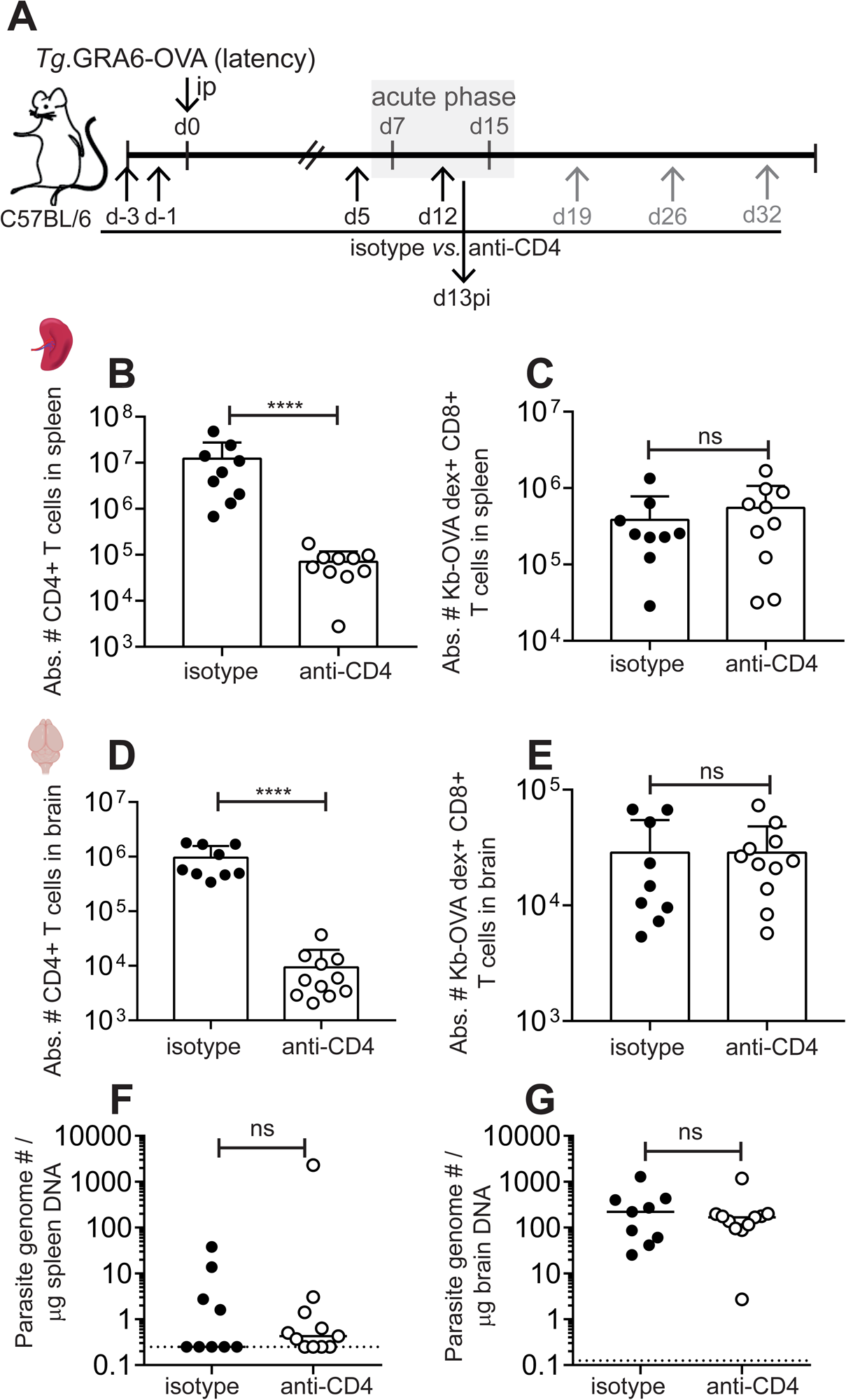
(related to Figure 4). CD4+ T cell depletion does not alter expansion of parasite-specific CD8+ T cells and parasite dissemination in spleen and brain during acute stage. (A) Schematics of experimental workflow: C57BL/6 mice were administered with an anti-CD4 depleting antibody at day -3 and -1 before infection with 200 tachyzoites of GRA6-OVA-expressing *T. gondii* Pru. Injection of anti-CD4 antibody was repeated at d5pi and d12pi until analyses of T cell responses and parasite burden at acute stage (d13pi). (B) Bar graph showing the absolute number of CD4+ T cells in spleen at d13pi. (C) Bar graph showing the absolute number of parasite (OVA)-specific CD8α+ T cells in spleen at d13pi. (D) Bar graph showing the absolute number of CD4+ T cells in brain at d13pi. (E) Bar graph showing the absolute number of parasite (OVA)-specific CD8+ T cells in brain at d13pi. (B, C, D, E) Each dot represents one mouse. Bars show the mean ± SD of N = 9 *vs*. 10 mice per group, pooled from 2 independent experiments with between 4 and 5 mice per group in each experiment. Mann-Whitney test between isotype-treated and anti-CD4-treated groups. (F, G) Parasite burden measured by qPCR on genomic DNA extracted from spleen (F) or brain (G). Each dot represents one mouse. Dotted line indicates the limit of quantification. Data pooled from 2 independent experiments with between 4 and 5 mice per group in each experiment. Mann-Whitney test between isotype-treated and anti-CD4-treated group.

**Sup. Figure 2.**
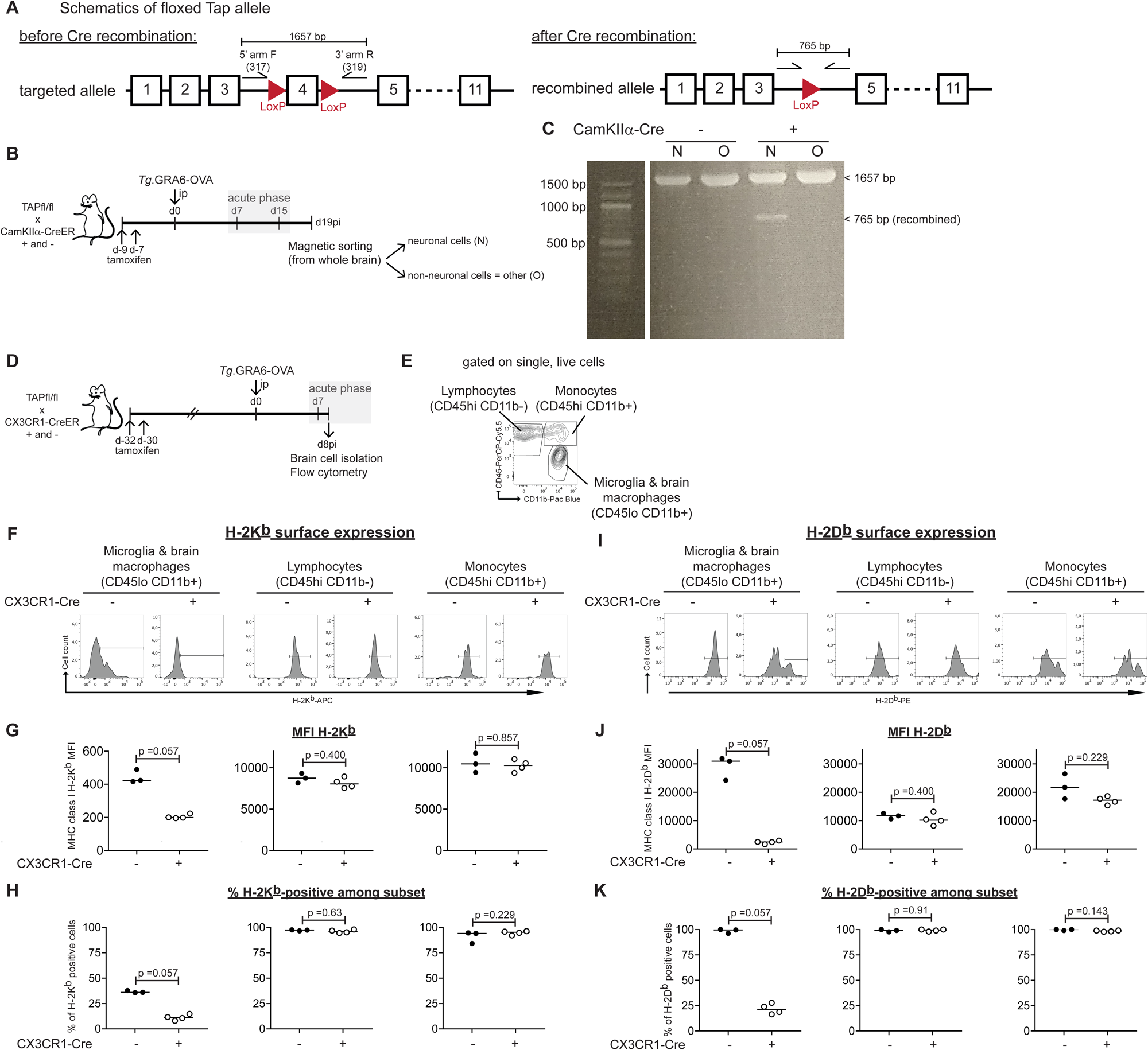
(related to Figure 5). Validation of Cre-mediated conditional invalidation of Tap1 gene in brain neurons and microglia/macrophages. (A) Schematics of floxed Tap1 allele: exon 4 of Tap1 gene (on chromosome 17) was flanked with 2 LoxP sites. PCR with 5’ arm F (#317) and 3’ arm R (#319) primers should produce a 1657-bp amplicon with the floxed (targeted) allele and a 765-bp amplicon with the post-Cre (recombined) allele. (B) Experimental workflow to validate Tap1 recombination in neurons following tamoxifen treatment. Tap1fl/fl X Camk2aCreER+ and – mice were treated with tamoxifen and infected one week later. At d19pi, neuronal (N) and non-neuronal cells (Other, O) were magnetically sorted from the whole brain, and genomic DNA was extracted. (C) PCR using 5’ arm F (#317) and 3’ arm R (#319) primers. The 765 bp-long amplicon corresponding to the Cre-recombined allele is detected only in neuronal cell fraction from tamoxifen-treated Camk2a-Cre+ mice. (D) Experimental workflow to validate Tap1 recombination in brain-resident microglia/macrophages by measuring MHC I expression by flow cytometry. Tap1fl/fl X Cx3cr1CreER+ and – mice were treated with tamoxifen and infected after one month, to allow renewal of the CX3CR1+ circulating monocytes as described in (Parkhurst et al., 2013). (E) At acute stage (d8pi), a phase during which inflammation and MHC I expression levels are maximal in the brain, mononuclear cells were isolated from the brain and stained with CD45 and CD11b to identify monocytes (CD45hi CD11b+), lymphocytes (CD45hi CD11b-), and brain-resident microglia and macrophages (CD45lo CD11b+). (F-K) Cells were stained with antibodies directed against H-2K^b^ and H-2D^b^ MHC I molecules. (F, I) Representative histograms depicting surface expression of H-2K^b^ (F) and H-2D^b^ (I) on CD45low CD11b+ brain-resident microglia and macrophages, CD45hi CD11b-lymphocytes and CD45hi CD11b+ monocytes. (G, H) Graphs showing the intensity of H-2K^b^ staining (G) and the proportion of H-2K^b^-positive cells (H) among the indicated sub-population. (J, K) Graphs showing the intensity of H-2D^b^ staining (J) and the proportion of H-2D^b^-positive cells (K) among the indicated sub-population. (G, J, H, K) Each dot represents one mouse. Data from one experiment. Cre+ and Cre-groups were compared with Mann-Whitney test.

**Sup. Figure 3.**
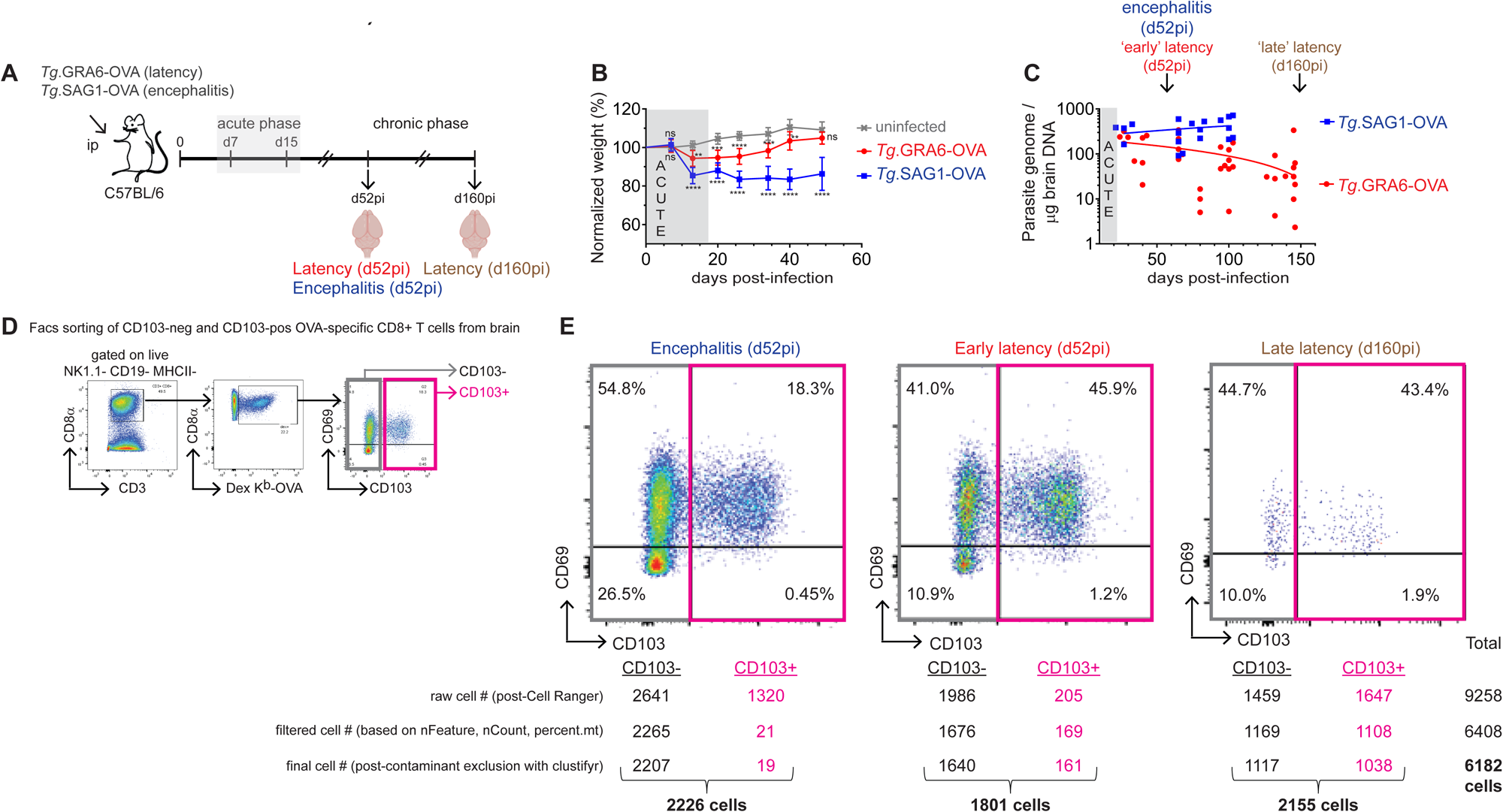
(related to Figure 6). Experimental workflow for single-cell RNA-seq analysis of parasite (OVA)-specific CD8+ T cells in encephalitis and latency infection models. (A) Schematics of experimental workflow: C57BL/6 mice were infected intra-peritoneally with 200 tachyzoites of GRA6-OVA- *vs*. SAG1-OVA-expressing *T. gondii* Pru, inducing respectively latency (red/brown) or encephalitis (blue). (B) Body weight monitored throughout infection, normalized with pre-infection value for each experimental group. Graph shows mean ± SD of N = 5 uninfected mice, N = 9 Tg.GRA6-OVA-infected mice, N = 8 Tg.SAG-OVA-infected mice. Two-way ANOVA with Tukey’s multiple comparison test applied on the 3 groups. Asterisks indicate statistical significance of the comparison between each infected group and the uninfected group. Results from one experiment representative of 3 independent experiments. (C) Brain parasite burden measured by qPCR on genomic DNA extracted from the brain. Each dot represents one mouse, results pooled from 4 experiments with Tg.SAG1-OVA-infected mice performed at 4 distinct timepoints and from 8 experiments with Tg.GRA6-OVA-infected mice performed at 8 distinct timepoints. (D) Gating strategy applied to Facs-sort CD103-negative and CD103-positive parasite (OVA)-specific CD8+ T cells isolated from the brain. A first gate was applied to select live, NK1.1-negative CD19-negative MHCII-negative cells, a second gate to select CD3+ CD8α+ T cells, a third gate to select OVA-specific CD8+ T cells thanks to SIINFEKL-loaded H-2K^b^ dextramers (dex K^b^-OVA). OVA-specific CD8+ T cells were ultimately separated between CD103-positive (all CD69+) and CD103-negative (including CD69+ and CD69-cells). (E) Flow cytometry dot plots showing CD69/CD103 labelings of Facs-sorted OVA-specific CD8+ T cells in the indicated conditions: early encephalitis (d52pi, 4 mice pooled), early latency (d52pi, 6 mice pooled) and late latency (d160pi, 10 mice pooled). In each condition, CD103-negative and CD103-positive cells were Facs-sorted and processed for scRNA-seq analysis. Percentages on the plots show the proportion of cells in each quadrant out of the parental OVA-specific CD8+ T cells. Numbers below the Facs plots indicate the cell number recovered for each category, following successive steps of quality control and contaminant exclusion performed with Seurat and clustifyr packages.

**Sup. Figure 4.**
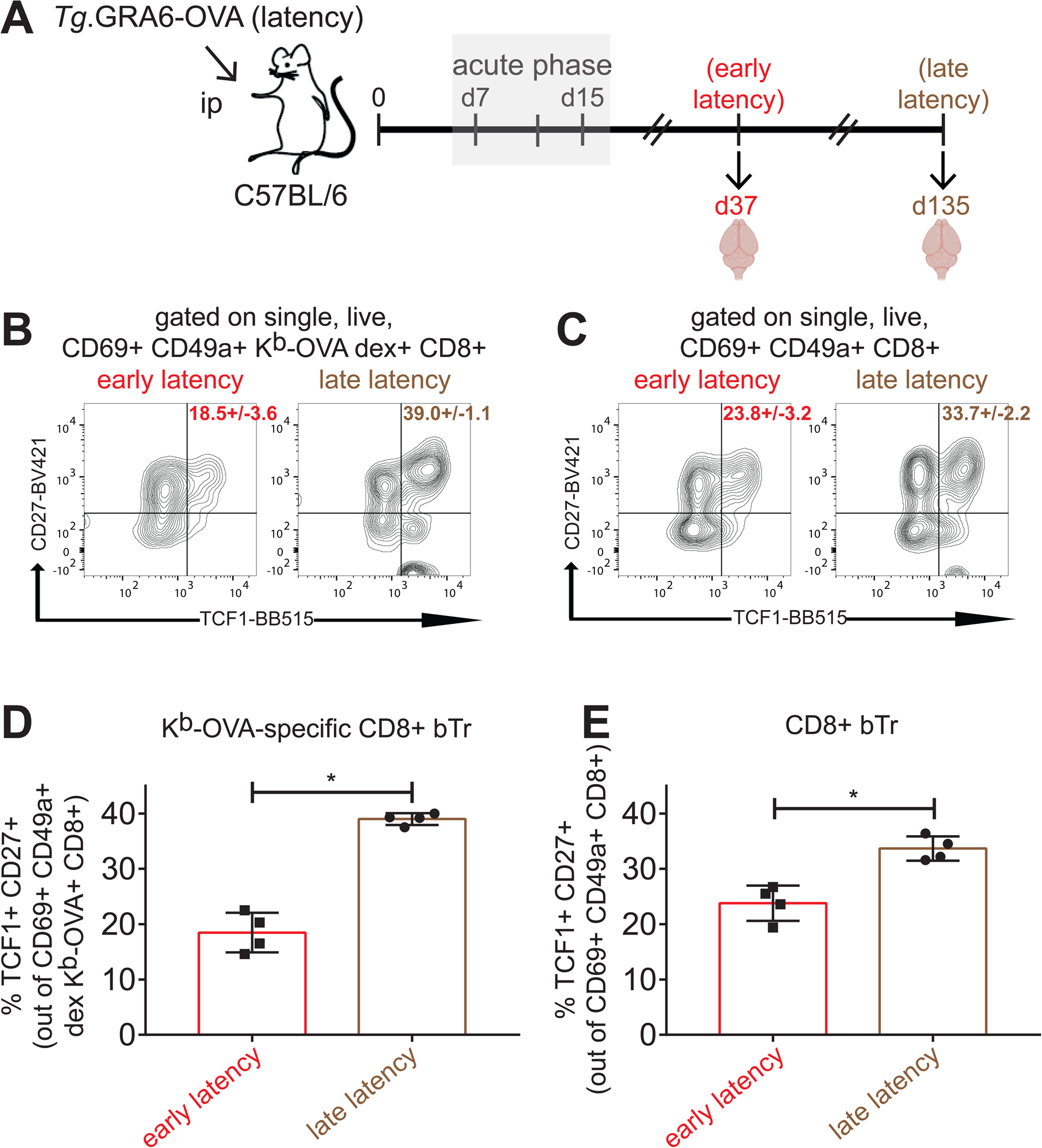
(related to Figure 6). A TCF1/CD27 double-positive ‘stem-like’ CD8+ bTr sub-population expands over time throughout *T. gondii* latent infection. (A) Schematics of experimental workflow: C57BL/6 mice were infected intra-peritoneally with 200 tachyzoites of GRA6-OVA-expressing *T. gondii* Pru, and analyzed at 2 time points during chronic stage: early (d37pi) and late latency (d135pi). (B, C) Facs plot of TCF1/CD27 stainings after gating on parasite (OVA)-specific CD8+ bTr (CD69+ CD49a+) (B) or total CD8+ bTr (CD69+ CD49a+) (C). Numbers on Facs plots show the percentage +/- s.d of the TCF1+ CD27+ subset out of OVA-specific (B) *vs.* total CD8+ (C) bTr cells. (D, E) Graph showing the percentage of TCF1+ CD27+ double-positive ‘stem-like’ cells out of parasite (OVA)-specific CD8+ bTr (D) or out of total CD8+ bTr (E). (D, E) Bars show mean ± s.d. of N = 4 mice, from one experiment. Mann-Whitney test between early and late latency group.

**Sup. Figure 5.**
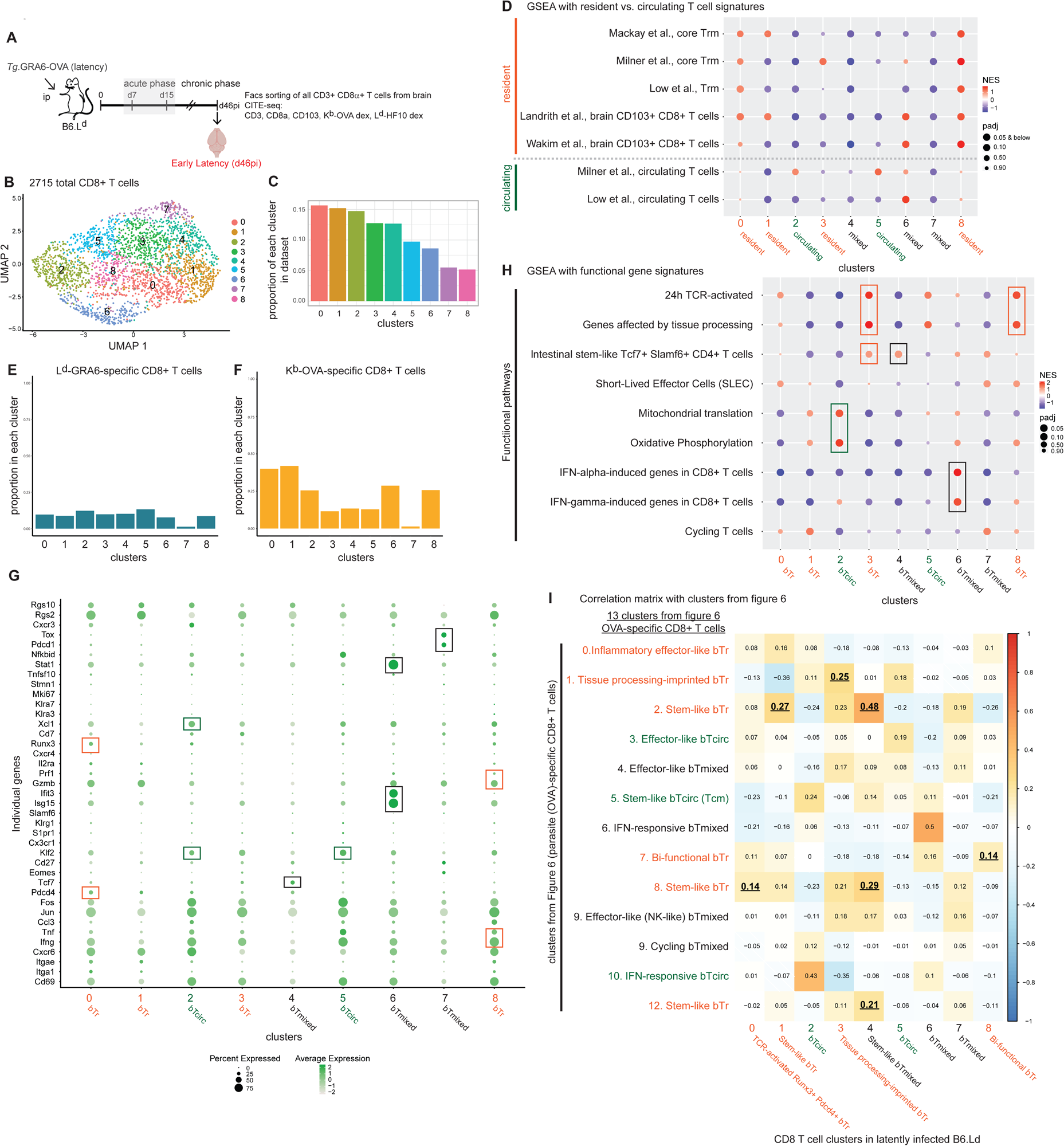
(related to Figure 6). CITE-seq analysis of brain-isolated total and *T. gondii*-specific CD8+ T cells upon latent infection of H-2L^d^-expressing mice. (A) Schematics of experimental workflow: H-2L^d^-transgenic C57BL/6 mice (B6.L^d^) were infected intra-peritoneally with 200 tachyzoites of GRA6-OVA-expressing *T. gondii* Pru. (B) Uniform Manifold Approximation and Projection (UMAP) plot of 2715 total CD8+ T cells isolated from the brain at early latency (d46pi, 4 mice pooled), partitioned in 9 clusters using Seurat-embedded Louvain clustering algorithm. (C) Bar graph showing the proportion of each cluster within entire dataset. (D) GSEA using tissue-resident T cell gene signatures (from (Mackay et al., 2013), (Milner et al., 2017), (Low et al., 2020), (Landrith et al., 2017), (Wakim et al., 2012)) *vs*. circulating T cell gene signatures (from (Milner et al., 2017) and (Low et al., 2020)). Each cluster is colored and tagged as ‘resident’ (bTr, orange) or ‘circulating’ (bTcirc, green), based on enrichment scores of resident *vs.* circulating T cell signatures. Clusters showing positive enrichment with both types of signatures (hybrid profile) were designated as ‘bTmixed’ (black-colored text). (E) Bar graph showing the proportion of L^d^-GRA6-specific CD8+ T cells, i.e. cells specific for the endogenous parasite antigen GRA6, among each cluster, as computed using CITE-seq with barcoded L^d^-GRA6 dextramers. (F) Bar graph showing the proportion of K^b^-OVA-specific CD8+ T cells among each cluster, as computed using CITE-seq with barcoded K^b^-OVA dextramers. (G) Dot plot showing average expression (color intensity) and percentage of gene-expressing cells (dot size) per cluster for a panel of selected genes. Boxes around the dots highlight a selection of upregulated genes of interest. (H) GSEA using previously published and/or public ‘functional’ gene signatures including: recent TCR activation (Low et al., 2020), imprinting of tissue dissociation procedure (van den Brink et al., 2017), stem-like CD4+ T cells (Schnell et al., 2021), short-lived effector cells (SLEC) (Joshi et al., 2007), mitochondrial translation (Reactome pathway knowledgebase (Gillespie et al., 2022)), oxidative phosphorylation (KEGG pathway database (Kanehisa et al., 2017)), type I and type II IFN responses (Mostafavi et al., 2016), proliferation (Clarke et al., 2019). (I) Correlation matrix between clusters of OVA-specific CD8+ T cells from infected C57BL/6 mice (see Fig. 6) and clusters of CD8+ T cells from infected B6.L^d^ mice, constructed using Hmisc package (rcorr function) to determine Spearman’s correlation between cluster pairs, taking into consideration the log2 fold-change of all genes per cluster.

